# Developmental Immunotoxicity of Low-dose Inorganic Arsenic Reprograms Macrophages Inducing Tumor-promoting Phenotypes

**DOI:** 10.1101/2025.07.14.663593

**Authors:** Emily J. Illingworth, Sylvia S. Sanchez, Kristal A. Rychlik, Andre Kleensang, Gabriel A. Lopez-Cecetaite, M. Carolina Rodriguez Steube, Aakriti Mathur, Pritam Sadhukhan, Daniel A. Medina-Cleghorn, Daniel Nomura, Martyn T. Smith, Mohammad O. Hoque, Fenna C.M. Sillé

**Affiliations:** Johns Hopkins University, Bloomberg School of Public Health, Department of Environmental Health and Engineering, Baltimore, MD, USA; Public Health Program, School of Health Professions, University of Mary Hardin-Baylor, Belton, Texas, USA; Department of Otolaryngology-Head and Neck Surgery, Johns Hopkins University School of Medicine, Baltimore, MD, USA; Department of Chemistry, University of California Berkeley, Berkeley, CA, USA; Department of Environmental Health Sciences, University of California Berkeley, Berkeley, CA, USA; Department of Oncology, Johns Hopkins University School of Medicine, Baltimore, MD, USA

**Keywords:** Inorganic arsenic, macrophages, metabolomic, lipidomics, cytokines, cancer

## Abstract

In many regions around the world, including the United States, inorganic arsenic (iAs) contaminates groundwater used for drinking, food production, and irrigation. Although the World Health Organization has set a safety limit of 10 µg/L for arsenic in drinking water, an estimated 200 million people worldwide are still exposed to arsenic concentrations above this threshold. Eliciting a broad range of adverse health effects, arsenic is a known carcinogen classified by the International Agency for Research on Cancer (IARC) and causes increased susceptibility to infectious diseases, highlighting its role as an immunotoxicant. The purpose of this study is to elucidate the effects of arsenic on the innate immune system, namely macrophages, using *in vitro* exposure models. Bone marrow-derived macrophages (BMDMs) were cultured from adult male and female C57/BL6 mice. These naïve macrophages (“M0” BMDMs) were exposed *in vitro* to a non-cytotoxic dose of iAs (0.1 µM sodium (meta)arsenite) during the 7 day period of macrophage differentiation and stimulated for 24 hrs with LPS and IFNγ (to induce “M1” pro-inflammatory activation) or IL-4 and IL-13 (to induce “M2” anti-inflammatory activation). In a parallel chronic exposure model, RAW 264.7 (RAW) macrophages were cultured *in vitro* with iAs for 70 days. Culture supernatant analysis for nitric oxide and cytokine secretion revealed sex-dependent differences in immune response between exposure models, as well as between iAs-exposed and nonexposed macrophages, with and without stimulation. Additionally, iAs-exposed macrophages exhibited increased lipid droplet formation and altered lipidomic and metabolomic profiles, as determined by LC/MS. Flow cytometric analysis further revealed changes in macrophage polarization markers in a sex- and stimulation-dependent manner, with M2-related markers being upregulated in iAs-exposed conditions. Finally, to assess the effects of iAs on macrophages in the context of cancer, we demonstrated that iAs-exposed macrophages displayed increased migration toward cancer cell-conditioned media, and promoted cancer cell proliferation. These results suggest that dysregulated macrophage polarization due to iAs exposure could impact susceptibility to diseases. This research contributes to our understanding of the full spectrum of adverse health effects of iAs exposure and may aid in the development of therapeutics for iAs-induced diseases, including cancer.

## Introduction

Inorganic arsenic (iAs) is a naturally occurring metalloid in the earth’s crust and remains an urgent public health concern due to its widespread contamination of drinking water and crop irrigation systems (WHO, 2006). Despite the World Health Organization (WHO) setting the drinking water safety standard of 10 µg/L (10 ppb), more than 200 million people worldwide continue to be exposed to arsenic at levels exceeding this standard (Podgorski & Berg, 2020; Shaji et al., 2021; WHO, 2006). In addition to being a potent carcinogen affecting the lung, skin, and bladder (IARC, 2012), both epidemiological studies (Banerjee et al., 2009; A. Rahman et al., 2011; Smith et al., 2011) and experimental findings support arsenic’s role as an immunotoxicant (Bishayi & Sengupta, 2006; Kozul et al., 2009; Lemaire et al., 2015; Lemarie et al., 2006, 2008; Medina et al., 2020; Sengupta & Bishayi, 2002). For example, individuals exposed to arsenic at levels above the WHO threshold show increased mortality from infectious diseases such as tuberculosis (Smith et al., 2011) and are more susceptible to lower respiratory tract infections (Rahman et al., 2011). Several reviews have broadly summarized evidence supporting arsenic’s role as an immunotoxicant (Bellamri et al., 2018; Dangleben et al., 2013) during pregnancy and early life (Attreed et al., 2017) and, specifically, on professional antigen presenting cells (Bahari & Salmani, 2017; Giles & Mann, 2022; Raqib et al., 2017, 2023).

Among these professional antigen-presenting cells, macrophages are a key target of arsenic toxicity. Macrophages are innate immune cells that recognize, phagocytose, process, and present antigens in the context of infectious disease and cancer (Hajishengallis & Russell, 2015; Hirayama et al., 2017; Parameswaran & Patial, 2010; Zhou et al., 2021). Macrophages are highly plastic and can polarize along a spectrum of activation (Mills et al., 2000; Orecchioni et al., 2019). For simplicity, we refer to these activation states as M0 (homeostatic or resting macrophage that has not seen a stimulus), M1 (“pro-inflammatory” macrophage, typically responding to infection), and M2 (“anti-inflammatory,” tissue remodeling macrophages). The activation states are determined by signals from their microenvironment (Mills et al., 2000; Orecchioni et al., 2019). For example, if macrophage toll-like receptor 4 (TLR4) is activated by lipopolysaccharide (LPS), a component of the outer membrane of gram-negative bacteria, this will trigger an M1 response characterized by pro-inflammatory cytokines such as TNF and IL-6 (Martinez & Gordon, 2014; Mosser & Edwards, 2008). Conversely, within the tumor microenvironment, for example, macrophages can be polarized toward an M2 phenotype by IL-10, IL-4, TGFβ, and metabolites of cancer cells like prostaglandins (Cui et al., 2017; Liu et al., 2021; Rőszer, 2015). These tumor-associated macrophages (TAMs) are often pro-tumorigenic and indicative of poor prognosis for many cancers due to their secretion of cytokines, such as IL-10 and TGFβ (Chen et al., 2019; Fiorentino et al., 1991; Zhang et al., 2016) and other metabolites (e.g., lipids) that drive tumor growth, angiogenesis, and promote an immunosuppressive microenvironment (Jayasingam et al., 2019; Liu et al., 2021).

Understanding how toxicants affect macrophage differentiation and polarization is critical for uncovering immune-mediated mechanisms of disease susceptibility. Disruption of the M1/M2 balance required for a normal immune response by environmental exposures, like arsenic, may compromise immune homeostasis. For example, Lemaire et al. (2011) demonstrated that arsenic-enhanced atherosclerosis is mediated by dysregulated lipid homeostasis of arsenic-exposed macrophages within atherosclerotic plaques (Lemaire et al., 2011). Further, in infectious disease models, decreased macrophage adhesion and chemotaxis were associated with reduced bacterial clearance of *Staphylococcus aureus* infection in mice (Bishayi & Sengupta, 2003). Finally, in the context of cancer, arsenic-exposed macrophages co-cultured with lung (Cui et al., 2017) or liver (Wang et al., 2021; Xue et al., 2021) cancer cells have been shown to enhance cancer cell proliferation compared to co-cultures with unexposed control macrophages. In some cases (Cui et al., 2017; Xue et al., 2021), this proliferative effect was accompanied by increased expression of M2 polarization markers, suggesting a shift toward an immunosuppressive phenotype. Arsenic also impacts the expression of genes related to hematopoiesis, which encompasses monocytes and myeloid-derived suppressor cells (MDSCs)--myeloid progenitors that are precursors to mature macrophages in the presence of macrophage colony-stimulating factor (M-CSF). These cell types play crucial, predominantly immunosuppressive, roles in the pathology of many arsenic-associated diseases, including cancers (Veglia et al., 2021).

Together, these data implicate macrophage differentiation and polarization as key mechanistic targets of arsenic exposure and underline the need for further characterization of arsenic-exposed macrophages in the context of various polarization stimuli. In this study, we utilized an *in vitro* exposure model of murine bone marrow-derived macrophages (BMDMs). BMDMs were exposed to noncytotoxic, human-relevant concentrations of inorganic arsenic to assess its effects on macrophage differentiation and polarization. We evaluated a range of M1- and M2-relevant endpoints, including nitric oxide (NO) production, differentiation and expression of M1- and M2-surfacemarkers via flow cytometry, cytokine secretion, lipid droplet formation, and comprehensive lipidomic and metabolomic profiling. Given the limited culture lifespan (∼1 week) of BMDMs, we also compared select findings in BMDMs to RAW 264.7 macrophages that were chronically exposed (70 days) to arsenic *in vitro* (Berghaus et al., 2010). Finally, to assess the effects of iAs on macrophages in the context of cancer, we determined whether iAs-exposed macrophages would increase migration toward cancer cell-conditioned media, and promote cancer cell proliferation.

## Methods

### Animals

Adult (8–10-week-old) male and female C57BL/6CR mice (five per sex) were purchased from Charles River Laboratories (Frederick, MD) and acclimated for 1 week prior to experimental procedures. Mice were cohoused five/cage and maintained in a temperature- and humidity-controlled facility on a 14:10 light:dark cycle and provided food and drinking water *ad libitum*. Mice received low arsenic chow (AIN-93M: Research Diets, New Brunswick, NJ, CAT# D10012Mi) and arsenic-free water (Crystal Geiser, Calistoga, CA).

### ARRIVE Statement

This study was conducted with approval by the Johns Hopkins University Institutional Animal Care and Use Committee (Protocols # MO19H465 and # MO20H283), following the National Research Council’s Guide for the Care and Use of Laboratory Animals. All animals were housed in the Johns Hopkins School of Public Health vivarium, which is in compliance with the Animal Welfare Act regulations and Public Health Service (PHS) Policy (PHS Animal Welfare Assurance #: D16-00173 (A3272-01)), and is accredited by the Association for the Assessment and Accreditation of Laboratory Animal Care (AAALAC) International (# 000503). In addition, we have been following the ARRIVE guidelines 2.0 (Percie du Sert et al., 2020) for the entire study.

### Isolation, Differentiation, and Polarization of Bone Marrow-derived Macrophages

Bone marrow from C57BL/6CR mice was isolated from the femurs and tibias of 8–10-week-old male and female mice, processed separately to maintain sex stratification for this study. Bones were flushed with phosphate buffered saline (1x PBS, pH 7.2, Gibco, Waltham, MA) containing 10% fetal bovine serum (FBS) using a 23G needle, and single cell suspensions were prepared using a 70 µm cell strainer. Red blood cells were lysed with ACK lysis buffer (Quality Biological, Gaithersburg, MD) and pelleted cells were resuspended and cultured in complete Dulbecco’s Modified Eagle Medium (DMEM) media containing 10% FBS, 200 µM L-glutamine, 100 µM HEPES (Gibco, Waltham, MA), 100 µM MEM Non-Essential Amino Acids (NEAA) (Gibco, Waltham, MA), 100 µM sodium pyruvate (Gibco, Waltham, MA), and 25 ng/mL of macrophage colony stimulating factor (M-CSF) (complete media) as previously described (Assouvie et al., 2018). Cells from each biological replicate were not pooled.

During the 7-day differentiation period, half the cells (∼2-4 x 10^7^) from each animal were maintained in media supplemented with inorganic arsenic (iAs) in the form of 0.1 µM sodium (meta) arsenite (Millipore Sigma, St. Louis, MO, CAT# S7400-100G). Cultures were maintained at 37 °C with 5% CO_2_ for 7 days with media refreshment (half of the media was replaced with 2x growth factors) +/- 0.1 µM sodium (meta) arsenite after 4 days. After 7 days of culture, complete and arsenic-containing media was removed and adherent cells were incubated with ice-cold PBS, harvested by manual cell scraping, and plated at 2-5 x 10^6^ cells per well in a clear, U-bottom 96-well plate. Cells were then polarized overnight to the M0 (complete medium alone), the M1 (complete media containing 300 ng/mL LPS and 6.25 ng/mL IFNγ), or the M2 (complete media containing 20 ng/mL each of IL-4 and IL-13) phenotype. Cells were not exposed to sodium (meta) arsenite during polarization. For lipidomics analysis, female M0 BMDMs were cultured as previously described, then treated with 1 µM monomethylarsonous acid (MMA(III)), a methylated metabolite of sodium arsenite, for 6 hours post-differentiation.

### Cell Line Cultures

For a chronic exposure macrophage model, RAW 264.7 macrophages (catalog TIB-71^™^, ATCC, Manassas, VA) were cultured in complete DMEM media as described above, without M-CSF, at 37°C with 5% CO_2_. Chronic exposure was modeled by culturing RAW macrophages with or without continual passage with 0.1 µM sodium (meta) arsenite for 70 days (12 passages).

For cancer cell cultures, we used lung (MLE-15, LLC1, KLN205) and bladder (MB49) tumorigenic cancer cell lines. MLE-15 tumorigenic lung cells (catalog T0495, Applied Biological Systems Inc., Waltham, MA) were cultured according to manufacturer’s protocol with RPMI medium with 2% FBS, 5 µg/mL insulin-transferrin-sodium selenite supplement, and 10 µM HEPES. LLC1 lung cancer cells (catalog CRL-1642, ATCC, Manassas, VA) and KLN205 cells (catalog CRL-1453, ATCC, Manassas, VA) were cultured according to manufacturer’s protocol with DMEM and 10% FBS. All cell lines were maintained at 37°C with 5% CO_2_.

### Cell Viability Assay

Cell viability was measured using a crystal violet staining assay as previously described (Feoktistova et al., 2016). BMDMs isolated from culture as described above, were seeded at 2 x 10^6^ cells in 200 µL volumes into flat-bottom 96-well plates and incubated overnight at 37°C with 5% CO_2_. The next day, plates were centrifuged at 1600 rpm for 10 minutes and the culture media was gently aspirated to remove non-adherent (presumed dead) cells. The remaining cells were washed once with PBS and adherent cells were fixed with 4% paraformaldehyde (PFA) for 10 minutes at room temperature. Once PFA was removed with aspiration, the cells were stained with 100 µL of 0.05% crystal violet solution for 30 minutes. Plates were then rinsed twice with tap water, drained, and dried inverted. The crystal violet stain was solubilized with 200 µL/well of methanol, the resultant supernatant was transferred to a new flat-bottom 96-well plate, and absorbance was measured at 570 nm using a microplate reader (Cytation 5, Biotek, Winooski, VT). Methanol-only wells served as blanks. Data are reported as fold-change in cell viability, comparing cancer cell proliferation in coculture with untreated versus arsenic-treated macrophages.

### Griess Assay

Nitric oxide (NO) was quantified in macrophage culture supernatant using the Griess assay (Amano & Noda, 1995). Briefly, 100 µL cell supernatant was collected from each well of stimulated macrophages in a new flat-bottom 96-well plate format. Griess reagents -- sulphanilamide and N-1-napthylethylenediamine dihydrochloride (NED) -- (50 µL each) were added consecutively to each well incubated at room temperature for 10 minutes. Absorbance was read at 540 using a microplate reader (Cytation 5, Biotek, Winooski, VT). Nitric oxide concentrations were determined using a sodium nitrite standard curve (with a range 0-200 µM NO) and expressed as µM of NO released in culture supernatant.

### Cytokine and Chemokine Multiplex Assay

The concentrations of 36 cytokines and chemokines were measured in macrophage culture supernatants using the R&D Systems Mouse Premixed Multi-Analyte Kits (LXSAMSM-30 and LXSAMSM-06). A complete list of all cytokines/chemokines measured is provided in **Supplemental Table 1**. Assays were performed following the manufacturer’s instructions and protocols optimized for use with the DropArray™ microplates and the Curiox plate washer system (Curiox, Woburn, MA) (Versteeg et al., 2017). All analytes were measured in technical duplicate and results were acquired using a Luminex MagPix system (Bio-Rad, Hercules, CA). Data was analyzed using xPONENT software. Cytokine concentrations were calculated from standard curves generated using five-parameter logistic regression and are reported in pg/mL.

### Flow Cytometric Analysis

Following the 7-day differentiation of bone marrow-derived macrophages, cells were suspended in phenol-free DMEM supplemented with 10% FBS, 200 µM L-glutamine, 100 µM HEPES, 100 µM MEM Non-Essential Amino Acids (NEAA), and 100 µM sodium pyruvate. Cells were counted using a hemocytometer with 0.4% trypan blue staining to determine cell viability. Viable cell counts were adjusted to a concentration of 2-10 x 10^6^ cells/ml and plated in a round-bottom 96-well plate. Macrophage polarization was performed as described above over a 24 hour period.

Post stimulations, cells were harvested by centrifugation at 1,600 rpm for 10 minutes at room temperature, supernatant was removed and cells were washed with 1X PBS. After another round of centrifugation and PBS removal, the cells were first stained with zombie aqua viability dye (Biolegend, San Diego, CA, CAT# 423102) according to manufacturer’s instructions. Upon removal of the Zombie dye buffer, the Fc receptors were blocked with antibodies (Biolegend, San Diego, CA, CAT# 101320) prior to surface staining. Cells were then stained for 30 minutes at 4°C with fluorochrome-conjugated antibodies targeting specific surface markers (see **Supplementary Table 4**). After surface staining, cells were washed, fixed and final preservation was performed in 1X BD FACS-lysing buffer (BD Biosciences, San Jose, CA, CAT# 349202) diluted in milli-Q ultrapure water (Milli-Q® IQ 7000 Ultrapure Lab Water System, MilliporeSigma, Burlington, MA). Intracellular staining was then performed to stain cytokines of interest using the BD Cytofix/Cytoperm (Catalog BD554714, BD Biosciences, San Jose, CA) protocol. Samples were individually run on a BD LSRII flow cytometer (Becton Dickinson, Franklin Lakes, NJ) utilizing FACSDiva software v6 and analyzed using FlowJo v10 (BD Biosciences, San Jose, CA). Compensation and gating strategies were applied using fluorescence minus one (FMO) and unstained controls.

### Lipid Droplet Assay

Lipid droplet formation was assessed using the Lipid Droplets Fluorescence Assay Kit (Cayman Chemical, Ann Arbor, MI, Item# 500001) (Listenberger & Brown, 2007) according to manufacturer’s instructions. Briefly, cultured macrophages were plated in a black, clear-flat-bottom 96-well plate at a density of 5 x 10^4^ cells/well in 200 µL volumes. As a positive control, the oleic acid supplement was added to culture media to a final dilution of 1:2000. The next day (∼18 hours later), cells were fixed and stained with 1 µg/mL Nile Red Solution. Relative fluorescence intensity was measured at FITC filter excitation/emission wavelengths of 485/535 nm on a microplate reader (Cytation 5, Biotek, Winooski, VT). Data are presented as relative fluorescence units (RFU), with background fluorescence subtracted from all readings.

### Fluorescent imaging of lipid droplets

After fluorescence intensity was quantified using the microplate reader, representative images of lipid droplets were acquired for each experimental condition using the R4 ECHO Revolve microscope (Discover Echo Inc., San Diego, CA). Stained cells were imaged directly in black, clear-flat-bottom 96-well formats, which was covered with a black box during imaging to minimize photobleaching from ambient light. A FITC filter cube (excitation/emission 485/535 nm) was used with a 10X objective lens and Ph1 phase ring condenser. Images were captured at a fixed exposure time of 565 ms.

### Metabolomics and Lipidomics Mass Spectrometry Analysis

For lipidomics analysis, BMDMs from female mice were cultured as described previously and treated with 1 µM monomethylarsonous acid (MMA(III)), a methylated metabolite of sodium arsenite, for 6 hours after differentiation. Macrophages (1 x 10^6^ macrophages) were pelleted, cell lysates were snap-frozen, and lipids were extracted using 40:40:20 acetonitrile:methanol:water for polar lipids (Kopp et al., 2010) or 2:1:1 chloroform:methanol:PBS for nonpolar lipids (Benjamin et al., 2013). Internal standards were included in all extractions. Lipid extracts were analyzed by LC/MS following established protocols.

For metabolomics analysis, BMDMs derived from both male and female mice were cultured with 0.1 µM iAs. Snap frozen cell pellets were resuspended in 50 µL of Milli-Q ultrapure water (Milli-Q® IQ 7000 Ultrapure Lab Water System, MilliporeSigma, Burlington, MA), washed with methanol, and extracted twice with methyl tert-butyl ether (MTBE), with speed vacuum drying between each extraction. Samples were vortexed, sonicated for 2 minutes, then centrifuged at 25,000 g for 30 minutes at 4°C. The lipid fraction (organic phase) was retained for future lipidomics comparison to MMA(III) treated samples. Metabolites were collected from the lower fraction (aqueous phase), subjected to another speed vacuum dry, then resuspended in 150 µL 1:1 acetonitrile:Milli-Q water for LC/MS analysis.

### Migration Toward Cancer Cell-conditioned Media

Macrophage migration in response to cancer cell secreted signals was assessed using a transwell migration assay. Conditioned medium was collected from cultures of murine lung carcinoma cells (LLC1 or KLN205) once they reached >85% confluency, as previously described (Guigni et al., 2020). To reduce potential contamination of any nonadherent cancer cells, the media was first centrifuged at 1,600 rpm for 10 minutes, and the supernatant was carefully removed without disturbing the pellet.

Migration assays were conducted using 24-well plates with 8.0 µm pore size transwell inserts (Corning, Corning, NY). Conditioned media (600uL) were added to the bottom wells, and BMDMs, cultured as described above, were seeded at 2 x 10^5^ per insert in 100uL of serum-free media. BMDMs were allowed to migrate overnight in a humidified incubator with 5% CO_2_, then the following day the media was aspirated from both top and bottom chambers. Cells were fixed with 4% paraformaldehyde (PFA) for 10 minutes at room temperature. Once PFA was removed with aspiration, the apical surface of the insert was swabbed with a cotton applicator to remove nonmigratory cells. The bottom of the insert was stained with 0.05% crystal violet solution for 30 minutes in the well to fully account for cells stuck to the basolateral transwell and those that migrated fully to the bottom. The inserts were rinsed twice with tap water, drained and dried inverted. The crystal violet stain was solubilized with 200 µL/transwell of methanol, and the methanol supernatant was transferred to a clean plate. Absorbance was measured at 570 nm using a microplate reader (Cytation 5, Biotek, Winooski, VT), with methanol-only wells as blanks. Results reflect relative migration efficacy and are presented as fold-changes compared to control conditions.

### Proliferation of Cancer Cells Cocultured with iAs-exposed Macrophages (MTT Assay)

To evaluate the impact of arsenic-exposed macrophages on cancer cell proliferation, indirect coculture experiments were conducted using 0.4 µm pore size transwell inserts (Corning, Corning, NY). Murine lung (MLE-15 and KLN205) and bladder (MB49) cancer cell lines were cultured, as previously described, and seeded at 1 × 10^4^ cells per well in the bottom chamber of the transwell plate in 600uL of complete mediim. In parallel, 1 × 10^5^ M0 BMDMs or RAW 264.7 macrophages that were previously exposed to iAs (in the form of 0.1 µM sodium (meta) arsenite), were seeded on the apical side of the transwells. The coculture was allowed to incubate for 6 days at 37°C with 5% CO_2_. After the coculture period, the transwell inserts were removed and the media was aspirated. To assess cell proliferation, a 300 µL volume of 1X MTT labeling reagent (diluted in PBS) was added to the bottom well. After an incubation of 24 hours, the MTT solution was solubilized with DMSO, and the plate was incubated for an additional 2 hours. Color change due to metabolic activity as a measure of proliferation of the cancer cells was measured using a microplate reader (Cytation 5, Biotek, Winooski, VT) at 550 nm using DMSO-only wells as blanks. Absorbance values were used as a proxy for cancer cell proliferation in response to macrophage derived signlaing. A subset of technical replicates of cocultured cancer cells were stored in DNA/RNA Shield (Catalog R1100, Zymo Research, Irvine, CA) for downstream RNA isolation and gene expression analysis.

### Gene Expression Analysis by Real-time Quantitative Polymerase Chain Reaction (RT-qPCR)

Total RNA was extracted from polarized murine bone marrow-derived macrophages using the Quick-RNA™ MiniPrep Kit (Zymo Research, Irvine, CA). RNA concentration was assessed using a NanoDrop™ 2000 spectrophotometer (Thermo Scientific™, Waltham, MA), and RNA quality was confirmed with the Ribogreen assay (Quant-it™ RiboGreen RNA Assay Kit, Invitrogen™, Waltham, MA). Complementary DNA (cDNA) was synthesized using the Applied Biosystems™ High-Capacity cDNA Reverse Transcription Kit (Thermo Scientific™, Waltham, MA), with total RNA diluted to a final concentration of 7.5 ng/μL.

RT-qPCR was performed in triplicate for all experimental groups--male- and female-derived BMDMs across each stimulation and arsenic treatment group--using the Applied Biosystems™ QuantStudio™ 3 Real-Time PCR System (Thermo Scientific™, Waltham, MA). Primers for target genes involved in macrophage polarization and metabolism were selected based on literature; sequences along with working annealing temps, are provided in **Supplemental Table 5**. Each 10 μL reaction included SsoFast™ EvaGreen^®^ Supermix (Bio-Rad Laboratories, Hercules, CA) and forward and reverse primers at 400 nM. Amplification curves were generated using an Applied Biosystems™ QuantStudio™ 3 System software (Thermo Scientific™, Waltham, MA) with the cycle conditions as follows: a hold stage (95°C for 2 minutes), 40 cycles at a PCR stage (95°C for 15 seconds, 56°C for 1 minute, 60°C for 1 minute), followed by a continuous melt curve analysis (95°C for 15 seconds, 60°C for 1 minute, and dissociation at 95°C for 15 seconds). Relative mRNA expression was calculated using the ΔΔCt method (Livak & Schmittgen, 2001), normalized to the housekeeping gene glyceraldehyde 3-phosphate dehydrogenase (GAPDH). Data were excluded if they did not meet the inclusion criteria of transcript detection at <33 cycles.

### Statistical Analysis

Nitric oxide production, lipid droplet content, and gene expression by RT-qPCR, were analyzed using unpaired Student’s t-tests. Multiplex cytokine, lipidomic, metabolomic, and flow cytometric data were analyzed using one-way ANOVA with Sidak’s multiple comparisons correction. When appropriate, analyses were stratified by sex, and results were normalized to cell viability. Unless otherwise indicated, significance is stated as compared to the sex-matched M0 untreated control. Statistical significance was defined as *p*-value <0.05. Statistical analyses were performed using GraphPad Prism 9 (Graphpad Software Inc, La Jolla, CA).

## Results

### Arsenic-induced Cytotoxicity to BMDMs

To establish a non-cytotoxic exposure dose for downstream analyses, we conducted a dose-response assessment of of sodium (meta) arsenite (0-1 µM) applied to BMDM cultures during the 7 day differentiation period. Briefly, to determine the cytotoxicity of iAs to BMDMs at each dose, we employed the crystal violet assay to measure cell viability. No significant cytotoxicity was observed at concentrations of 0.0001, 0.001, 0.01, or 0.1 µM iAs doses; however, exposure to 1 µM resulted in a significant ∼18% reduction in viability compared to the control (**Fig. 1**). To avoid off-target effects of BMDMs responding to significant amounts of dead and/or dying cells, we proceeded with the highest non-cytotoxic concentration of iAs, 0.1 µM, for subsequent experiments. It is important to note that this concentration equates to ∼13 ppb, slightly above the WHO drinking water guideline of 10 µg/L, and is therefore relevant to represent human exposure levels (WHO, 2006).

**Figure 1.**
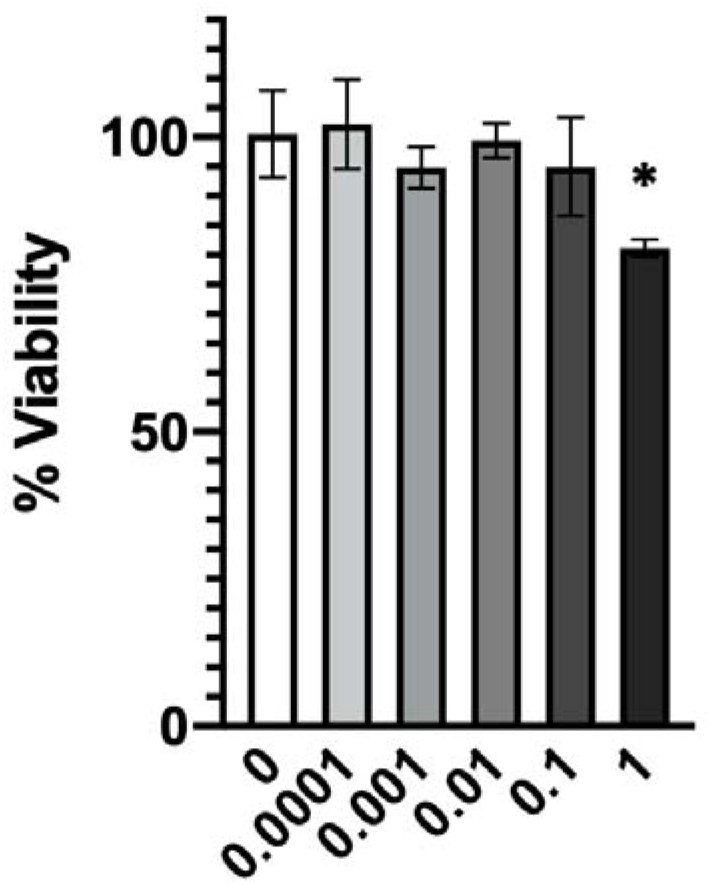
Cytotoxicity of iAs to BMDMs. M0 BMDMs were treated with a dose range of iAs (0-1 µM) and viability was measured by crystal violet staining and absorbance. Plots shows mean ± SD with % viability normalized to nonexposed macrophages and dose range across the x-axis. \**p*-value<0.05. iAs=inorganic arsenic; BMDMs=bone marrow-derived macrophages.

### iAs Exposure Reduces Nitric Oxide Production in iAs-treated Macrophages

Macrophage nitric oxide (NO) production is a hallmark of M1 polarization in response to inflammatory signals, including cytokines and/or microbial pathogen-associated molecular patterns (PAMPs) (Tripathi, 2007). Since this is an important M1 endpoint, we evaluated NO production in culture supernatant from both iAs-exposed BMDMs and RAW 264.7 macrophages using the Griess assay. No significant differences in NO production were observed in either male- or female-derived BMDMs exposed to 0.1 µM iAs during differentiation following stimulation with 300 ng/mL LPS plus 6.25 ng/mL IFNγ (**Fig. 2A, 2B**). However, RAW 265.7 macrophages chronically exposed to the same dose of arsenic exhibited a significant reduction in NO release following M1 stimulation (*p<0.05*; **Fig. 2C**). As expected and previously known (Mills, 2001), NO production was negligible in both M0- and M2-stimulated BMDMs and RAW 264.7 macrophages (data not shown).

**Figure 2.**
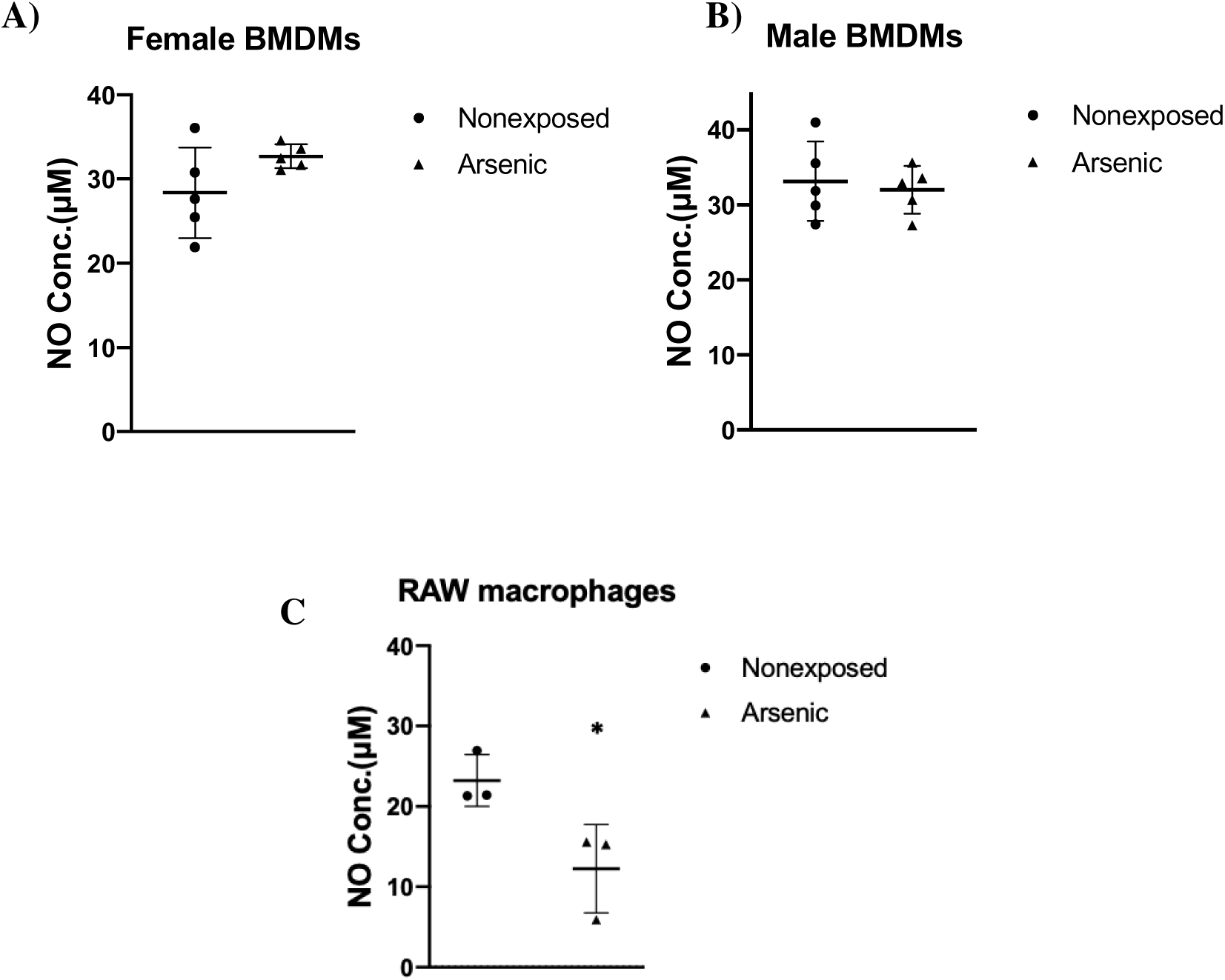
NO release by iAs-exposed M1 macrophages. NO release was measured by the Griess assay on culture supernatant obtained from female (A) or male (B) M1-stimulated BMDMs exposed to 0.1 µM iAs for 1 wk during differentiation, and from M1-stimulated RAW 264.7 macrophages (C) exposed to iAs for 70 d. n=5 biological replicates for BMDMs and n=3 technical replicates for RAW 264.7 macrophages. M0- and M2-stimulated macrophages from both culture models had negligible production of NO (data not shown). Plots show mean ± SD. \**p*-value <0.05. NO=nitric oxide; iAs=inorganic arsenic; BMDMs=bone marrow-derived macrophages.

### Cytokine and Chemokine Profiles of iAs-treated BMDMs

In response to signals they receive from their microenvironment, macrophages produce cytokines and chemokines that signal in autocrine, paracrine, and endocrine fashion to help orchestrate an immune response (Mantovani et al., 2004; Rostam et al., 2017). To determine the effects of arsenic exposure on cytokine profiles of M0-, M1-, and M2-stimulated macrophages, we performed multiplex cytokine and chemokine analysis on cell supernatant harvested from BMDMs 24 hours post-stimulation. A total of 36 analytes were measured and we found significant changes in cytokine/chemokine profiles of iAs-treated BMDMs that were sex- and stimulation-dependent. Female M0 (unstimulated) BMDMs that were iAs-treated showed a decrease in TIMP-1 secretion (*p*<0.01) and an increase, although not statistically significant (*p*=0.14), in CCL5, compared to unexposed matched control BMDMs, whereas iAs-treated male M0 (unstimulated) BMDMs had a significant reduction in CHI3L1 and CXCL2 (*p* <0.001 and *p* <0.0001, respectively), compared to unexposed male control BMDMs (**Fig. 3A, 3B**). Under M1 stimulation, iAs exposure significantly decreased CCL5 in females (p<0.05) and CCL4 in males (p<0.0001). A similar but nonsignificant trend was oberseved in CCL4 signaling in female BMDMs (p=0.053) (**Fig. 3C, 3D**). Male and female M2-stimulated BMDMs that were iAs-treated similarly showed a decrease in chemokine secretion, with iAs-treated females demonstrating significantly decreased expression of CCL4 (*p*<0.01), CCL5 *(p*<0.001) and CXCL2 *(p*<0.001), while iAs-treated males showed significantly reduced CXCL2 expression (*p*<0.001) compared to untreated controls (**Fig. 3E, 3F**). Collectively, these data demonstrate that iAs exposure suppresses chemokine (CCL4, CCL5, and CXCL2) and cytokine (CHI3L1 and TIMP-1) secretion in both male and female macrohphages across stimulation conditions.

**Figure 3.**
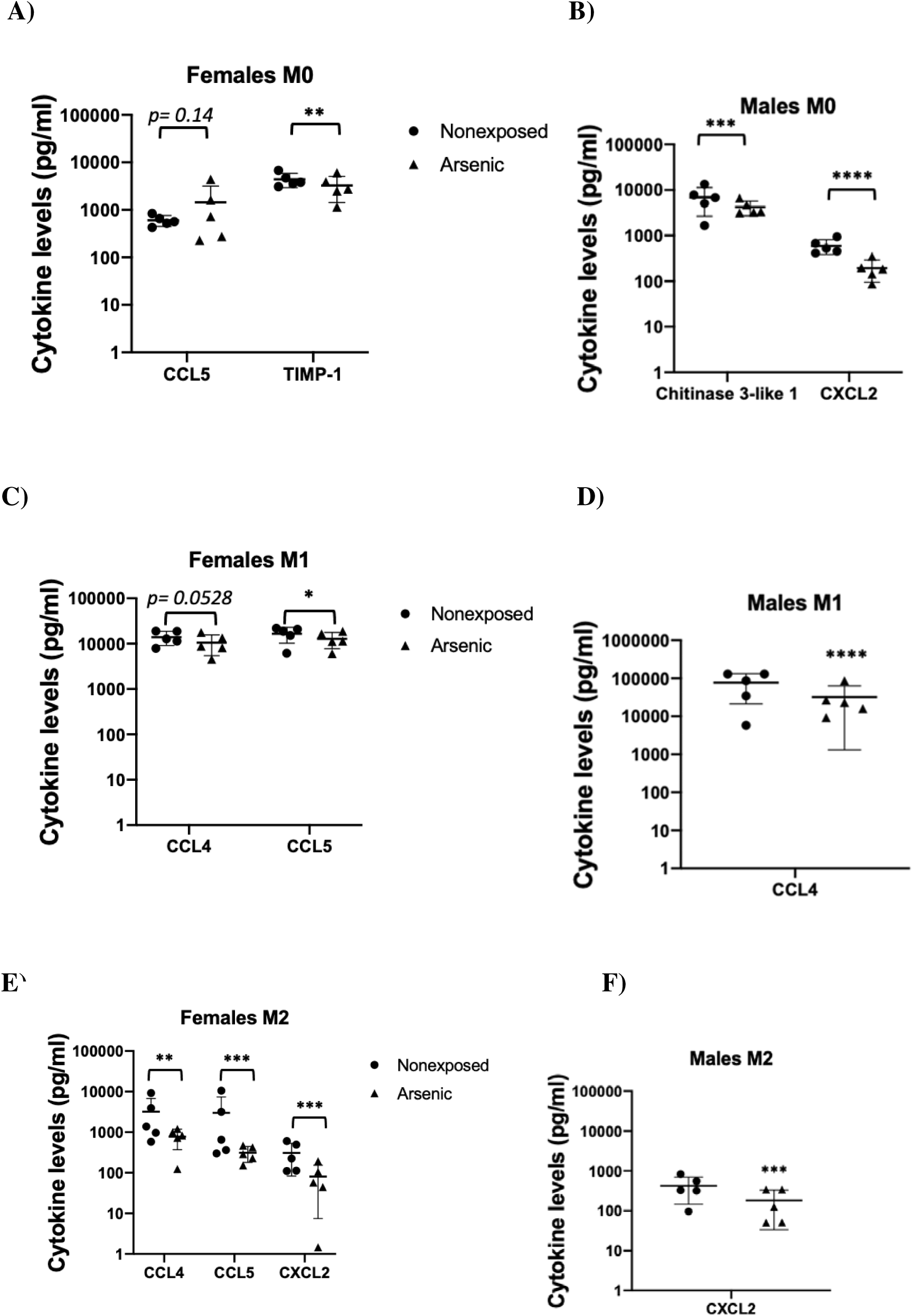
Changes in cytokine profiles of iAs-exposed macrophages. Using a multiplex assay, cytokine concentrations for 36 analytes were measured from culture supernatant of M0-(A, B), M1-(C, D), or M2-stimulated (E, F) female and male BMDMs exposed to 0.1 µM iAs for 1 wk during differentiation. n=5 biological replicates. Only analytes reaching or approaching significance are shown. Plots show mean ± SD. \**p*-value<0.05; \*\**p*-value<0.01; \*\*\**p*-value<0.001; \*\*\*\**p*-value<0.0001. BMDMs=bone marrow-derived macrophages; iAs=inorganic arsenic.

### Flow Cytometric Characterization of iAs-treated BMDMs

To assess the effects of arsenic exposure on macrophage differentiation and polarization, we designed a multicolor flow cytometry panel (**Supplemental Table 4**) including markers of monocyte-macrophage differentiation (Ly6C, CD11b, F4/80), polarization status (iNOS, Arg1, CD163), and functional capacity (CD86, MHCII, PD-L1, TGFβ, IL-6) across M0-, M1-or M2-polarization. For these studies, we defined mature macrophages as CD11b^+^F4/80^+^ (Zajd et al., 2020), while CD11b^+^ Ly6C^+^ F4/80^-^MHCII^-^ cells were included to capture monocytic precursors and myeloid-derived suppressor cells (MDSCs) (Bronte et al., 2016; Damuzzo et al., 2015) (for a representative gating strategy, see Supplemental **Fig. 3**).

In female BMDMs, arsenic exposure during differentiation led to a nonsignificant increase in Arg1-expressing mature macrophages after M2 stimulation (*p*= 0.119, **Fig. 4A**). No significant changes in monocyte to macrophage differentiation were observed in either male-or female-derived BMDMs as evidenced by similar percents of mature macrophages (CD11b+F4/80+), although the iAs-exposed male-derived M2 group showed a slightly reduced percentage of mature macrophages compared to… (*p*=0.111, **Fig. 4D**). TGFβ production was decreased nonsignificantly in iAs-treated M0 male-derived BMDMs (*p*= 0.127, **Fig. 4C**), while CD163 expression increased in M1 stimulated iAs treated male BMDMs, though this trend was also not statically signicant (*p*= 0.120, **Fig. 4B**).

**Figure 4.**
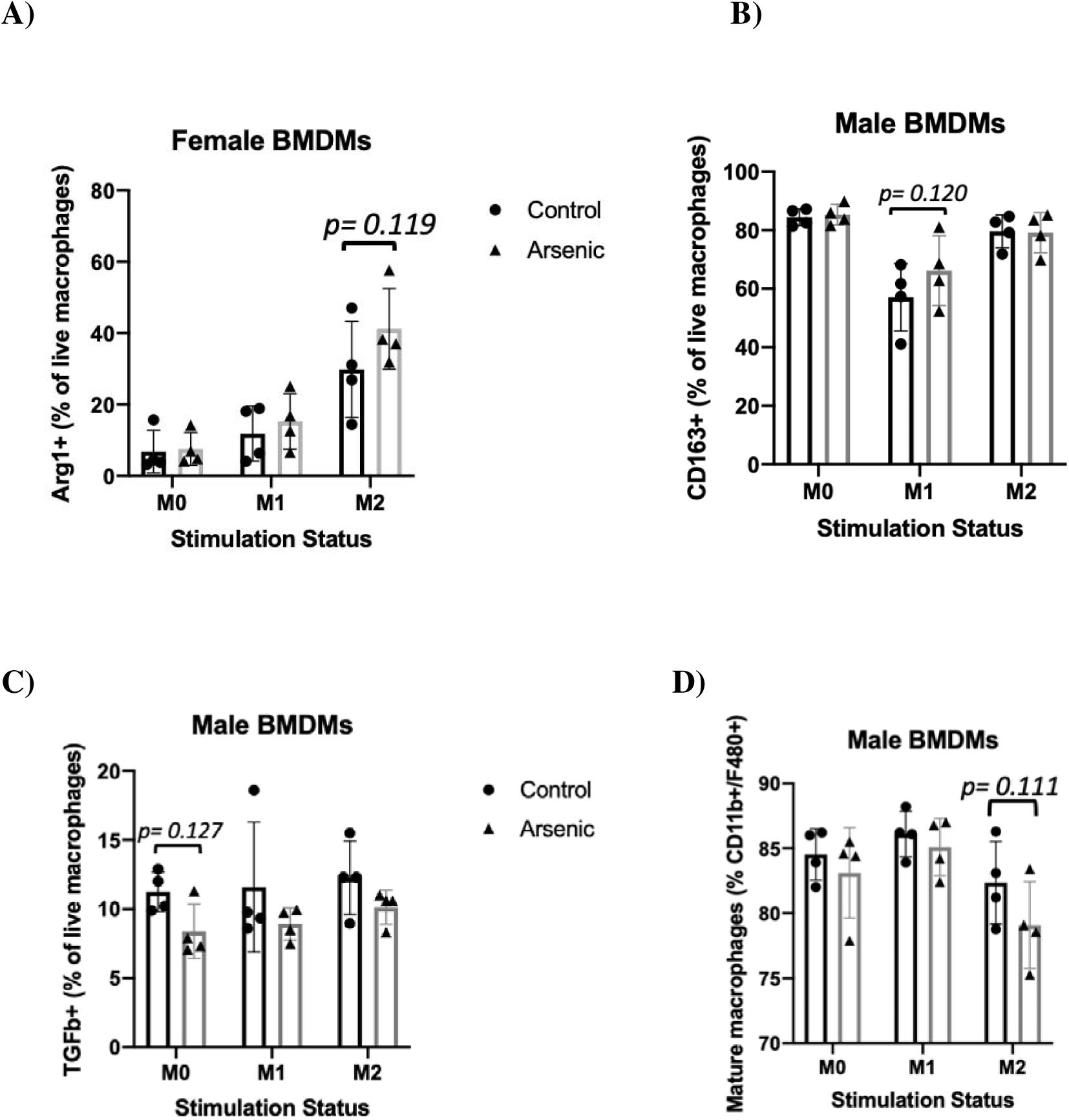
Flow cytometric characterization of iAs-exposed female and male BMDM polarization and functional markers as a frequency of parent gates. Mature macrophages (CD11b+F4/80+ live cells) were analyzed from BMDM cultures for expression of select polarization and functional markers Arg1 (A), CD163 (B), TGFβ (C), and for culture homogeneity as a percent of live cells (D). Only plots with *p*-values <0.15 are included (complete panel with all markers analyzed can be found in Supplemental Table 4). Plots show % expression of parent gate on the y-axis by stimulation condition on the x-axis, with the mean ± SD. n=4 biological replicates. \**p*-value<0.05. iAs=inorganic arsenic; BMDMs=bone marrow-derived macrophages

Among mature macrophages (CD11b+F4/80+), the level of protein production (reflected as mean fluorescent intensity, or MFI) was altered for a variety of markers. In iAs-treated M2-stimulated female- and male-derived BMDMs, Arg1 expression was increased, though only significantly in males (*p*= 0.109 and *p*<0.05, respectively, **Fig. 5A, 5C**). CD86 followed the same trend of upregulation with only the male-derived group reaching significance (*p= 0.113* and *p*<0.05, for females and males, respectively, **Fig. 5B, 5D**). In contrast, the expression of iNOS was significantly reduced in M1-stimulated male BMDMs (*p*<0.01, **Fig. 5E**) and the expression of MHCII was decreased in both M1 and M2 groups (p=0.056 and *p*<0.01, respectively, **Fig. 5F**).

**Figure 5.**
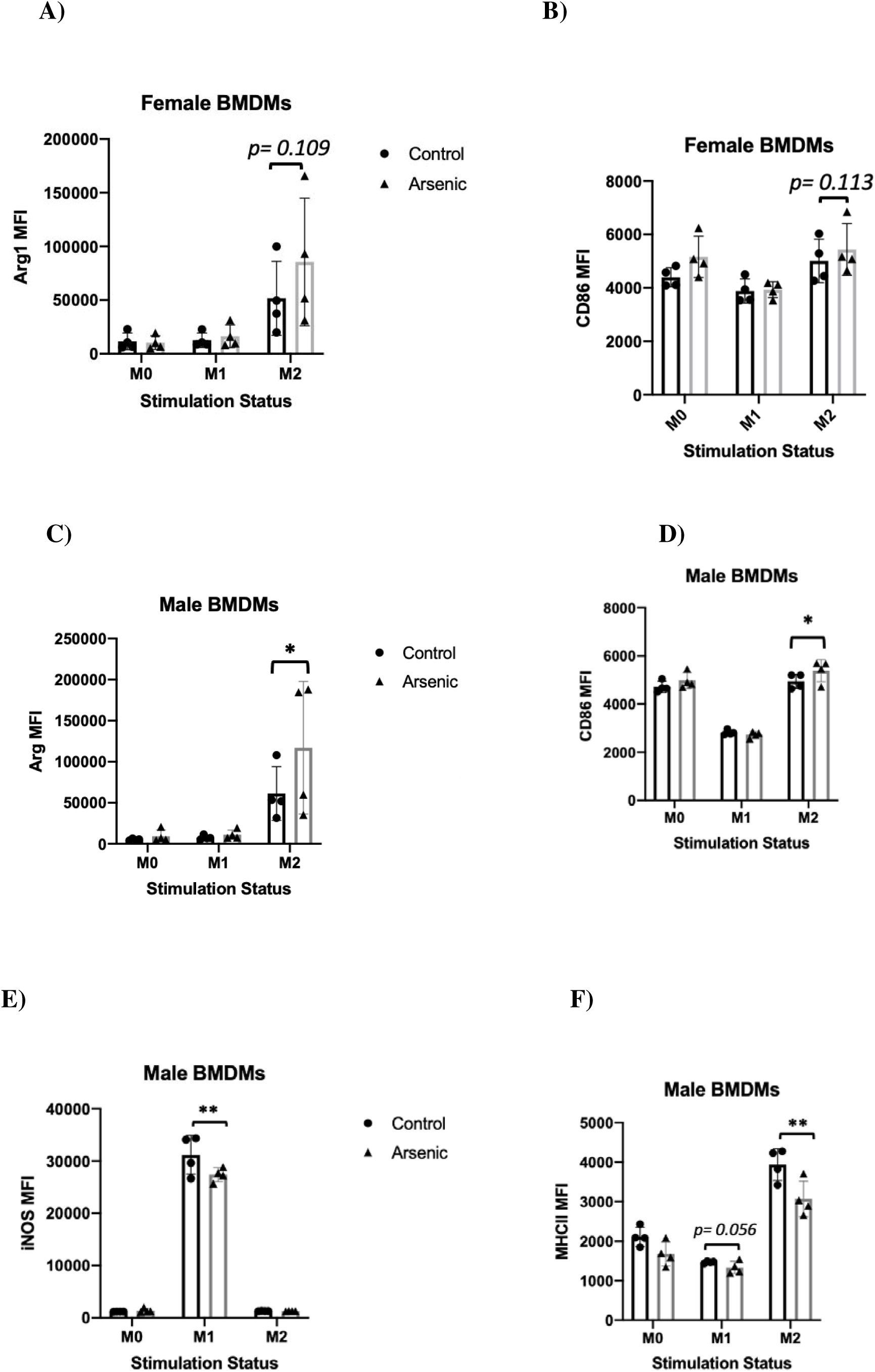
Flow cytometric characterization of iAs-exposed female and male BMDM polarization and functional markers as MFI expression levels. Mature macrophages (CD11b+F4/80+ live cells) were analyzed from BMDM cultures for expression of select polarization and functional markers Arg1 (A, C), CD86 (B, D), iNOS (E), and MHCII (F). Only plots with *p*-values <0.15 are included (complete panel with all markers analyzed can be found in Supplemental Table 4). Plots show MFI on the y-axis by stimulation condition on the x-axis, with the mean ± SD. n=4 biological replicates. \**p*-value<0.05; \*\**p*-value<0.01. iAs=inorganic arsenic; BMDMs=bone marrow-derived macrophages. MFI= mean fluorescence intensity.

There were no differences in the frequency of MDSCs in our iAs-exposed cultures, though differences were observed for frequency and expression levels of certain markers. The frequency of CD163-expressing MDSCs was increased in the iAs-treated female M1-stimulated group and the iAs-treated male M0 group upon iAs exposure, though only the females reached statistical significance (*p*<0.05 and *p*= 0.102, respectively, **Supp. Fig. 1A, 1B**). Similar to our findings in BMDMs, the frequency of live MDSCs expressing iNOS was significantly reduced in the iAs-treated male M1-stimulated group (*p*<0.05, **Supp. Fig. 1D**) and CD86 expression trended lower in the same group (*p*= 0.122, **Supp. Fig. 1C**). Male M2-stimulated MDSCs with iAs treatment had a nonsignificant increase in the frequency of PD-L1 expression compared to controls (*p*= 0.079, **Supp. Fig. 1E**). iAs exposure also affected the MFI of certain markers on MDSCs. CD86 expression decreased in M1-stimulated cells in both females and males, though not significantly (*p*= 0.107 and *p*= 0.121, respectively, **Supp. Fig 2B, 2C**). In M2 female MDSCs only, iAs significantly increased the MFI of CD163 (*p*<0.05, **Supp. Fig. 2A**). In M1 male MDSCs only, iAs significantly increased the MFI of PD-L1 (*p*<0.05, **Supp. Fig. 2D**). As shown by the decreased expression of pro-inflammatory M1 markers and the increased expression of M2 markers in arsenic-treated BMDM cultures across stimulation groups, these data collectively suggest that chronic arsenic exposure skews macrophages and MDSCs toward the M2 phenotype.

### Lipid Droplet Formation in iAs-treated BMDMs and RAW 264.7 Macrophages

Macrophages have different modes of lipid metabolism based on their polarization state. M1 macrophages favor aerobic glycolysis and fatty acid synthesis, while M2 macrophages rely on fatty acid oxidation and oxidative phosphorylation (Kelly & O’Neill, 2015; Morgan et al., 2021; Rodríguez et al., 2025). Because of the metabolic reprogramming that coincides with macrophage polarization, we investigated whether arsenic exposure skewed the metabolic signature of M0 (homeostatic) macrophages. First, we assessed whether iAs treatment affected the formation of lipid droplets in either M0 BMDMs or M0 RAW 264.7 macrophages. Briefly, BMDMs or RAW 264.7 macrophages were cultured, plated, and stained with Nile Red, a lipid indicator. Fluorescence intensity was measured, and representative images were captured using the ECHO Revolve fluorescence microscope. No significant differences were observed between iAs-treated and control male or female BMDMs (**Fig. 6A, 6B**). However, chronic iAs exposure significantly increased lipid droplet accumulation in M0 RAW 264.7 cells compared to controls (**Fig. 6C, 6D**). These findings demonstrate that chronic arsenic treatment enhances lipid droplet biogenesis in M0 RAW 264.7 macrophages.

**Figure 6.**
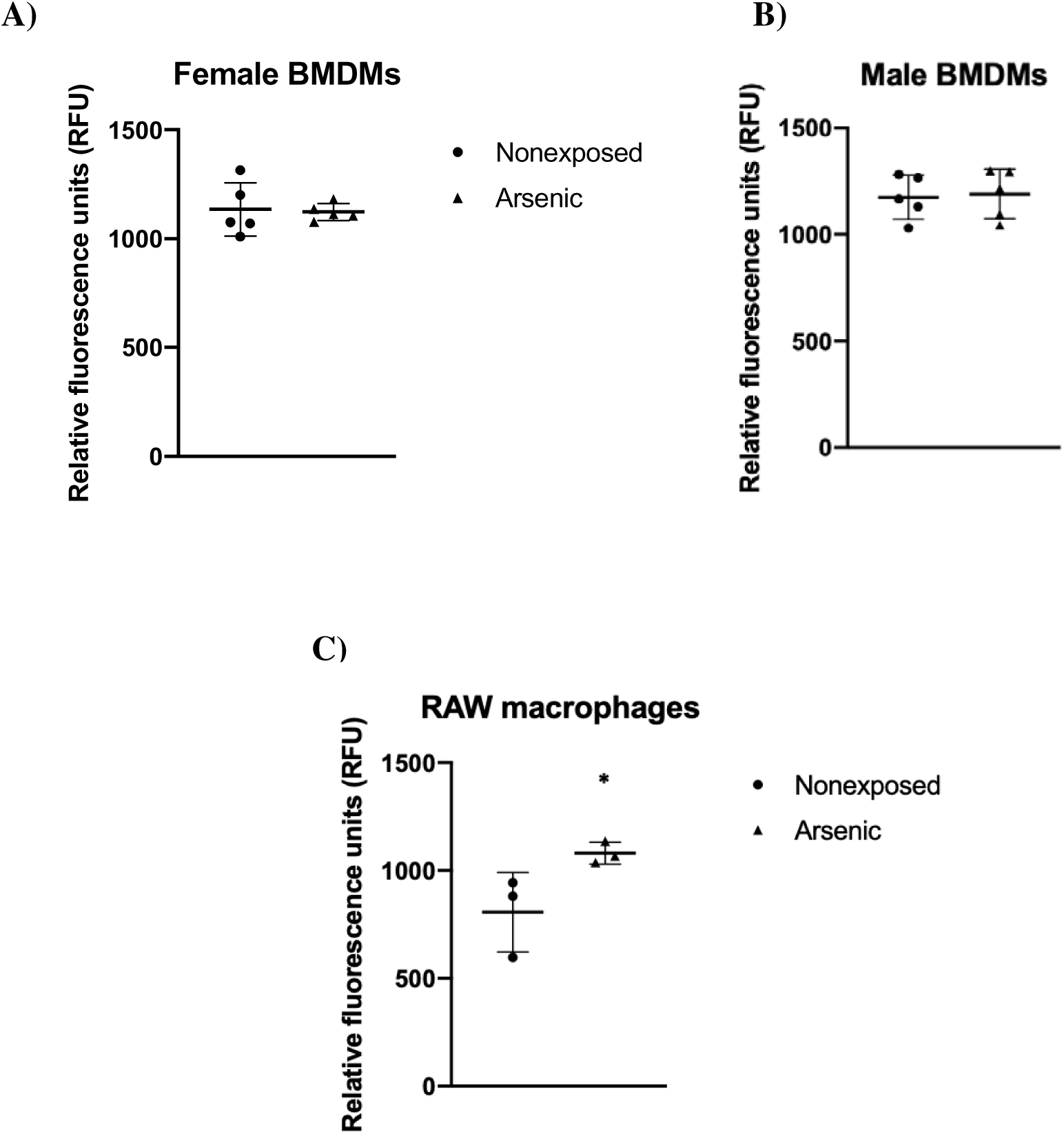
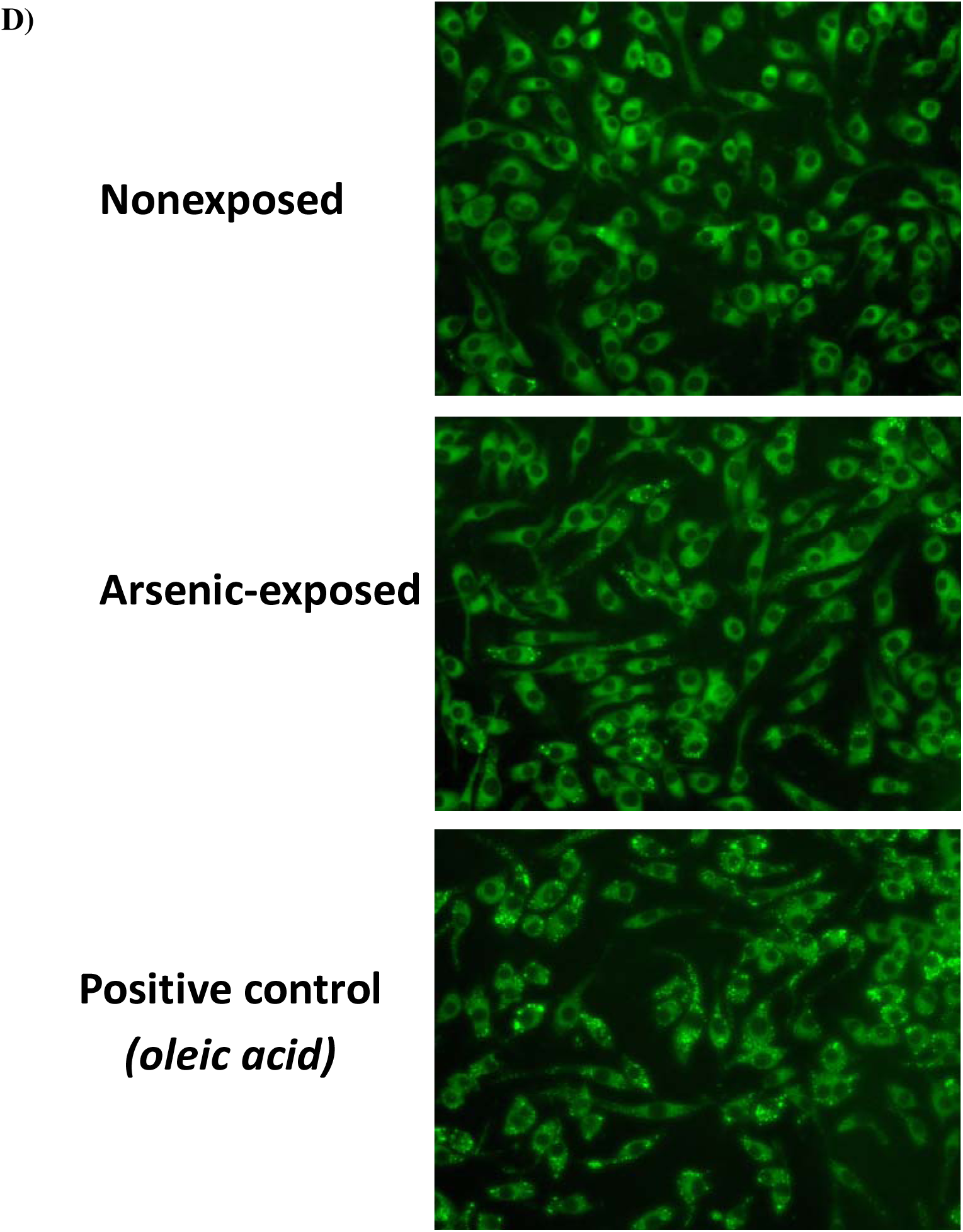
Lipid droplet formation in iAs-exposed M0 macrophages. Lipid droplet formation was measured using a lipid droplet fluorescence assay in female (A) or male (B) BMDMs exposed to 0.1 µM iAs for 1 wk during differentiation, and from RAW 264.7 macrophages (C) exposed to iAs for 70 days from which representative images are shown (D). n=5 biological replicates for BMDMs and n=3 technical replicates for RAW 264.7 macrophages. Plots show mean ± SD. *p-value<0.05. iAs=inorganic arsenic; BMDMs=bone marrow-derived macrophages.

### Lipidomic Profiles of iAs-treated BMDMs

Since increased lipid droplet formation was observed in chronic iAs-exposed M0 RAW 264.7 macrophages (**Fig. 6C, 6D**), it was of particular interest to detect any alterations in the levels of lipid metabolites of iAs-treated macrophages using a more sensitive method. Female BMDMs were M0 stimulated and cultured as previously described, then treated with monomethylarsonous acid (MMA(III)), a methylated metabolite of sodium arsenite, for 6 hours after differentiation. M0 macrophages were pelleted, snap frozen, and polar and nonpolar lipids were extracted (Benjamin et al., 2013; Kopp et al., 2010) and analyzed by LC/MS. Of the 30 lipids measured across 6 lipid classes (**Supplemental Table 2**), 11 were significantly altered following MMA(III) exposure (**Fig. 7**). Acylcarnitines were dysregulated with arsenic exposure, with lauroyl carnitine (C12:0) significantly downregulated *(p*<0.0001) while palmitoyl L-carnitine (C16:0) showed a trending but nonsignificant upregulation (*p*= 0.0968). Other lipid targets that were significantly altered and upregulated by arsenic exposure were as follows (lipid classes bolded): **eicosanoids** prostaglandin E2 and D2 (PGE2/PGD2, *p*<0.0001), **ether lipids** lysophosphatidic acid-platelet activating factor (lyso-PAF [C18:1], *p*<0.0001) and phosphatidic acid ether (PAe [C16:0/C20:4], *p*<0.001), **lysophospholipids** lysophosphatidic acid (LPA [C20:4], *p*<0.0001), lysophosphatidylcholine (LPC [C20:4], *p*<0.01), lysophosphoethanolamines (LPE [C18:1], *p*<0.0001; LPE [C18:0], *p*<0.0001; LPE [C20:4], *p*<0.05; LPS [C18:0], *p*<0.05) and one analyte from **the sphingolipids,** sphingosine 1-phosphate (S1P [C16:0], *p*<0.001) (**Fig. 7**). These results suggest that female-derived BMDMs exposed to MMA(III), a toxic metabolite of sodium (meta) arsenite, have enhanced production of lipid mediators associated with inflammation and tumorigenesis. In the future, we will repeat this analysis with sex- and stimulation-stratified BMDMs treated with sodium (meta) arsenite.

**Figure 7.**
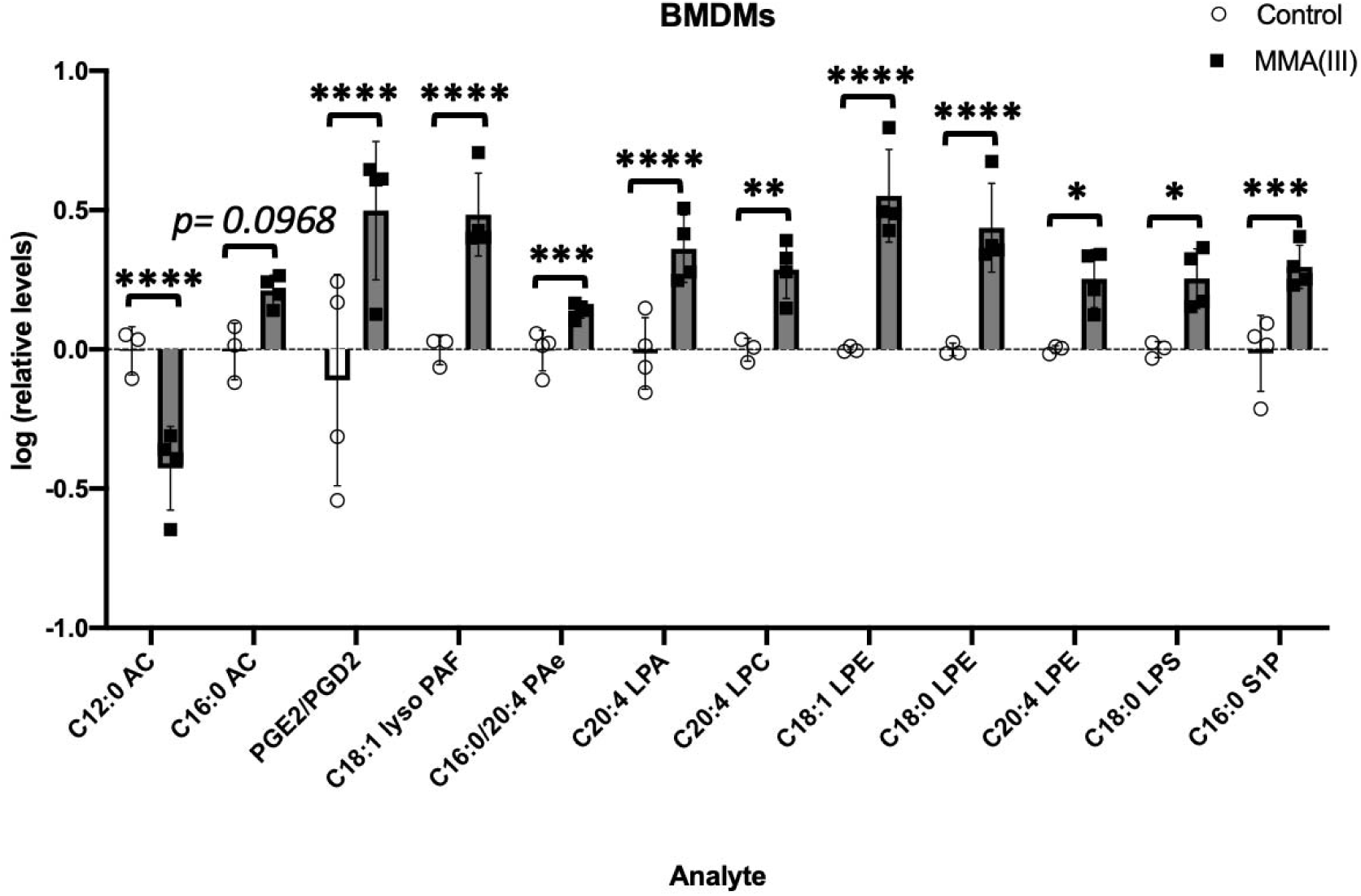
Lipid profiles of MMA(III)-exposed female BMDMs. Polar and nonpolar relative lipid levels from BMDMs exposed to 1 µM MMA(III) for 6 hrs were determined by LC/MS. n=3-4 biological replicates. Plots show mean ± SD. \**p*-value<0.05; \*\**p*-value<0.01; \*\*\**p*-value<0.001; \*\*\*\**p*-value<0.0001. iAs=inorganic arsenic; BMDMs=bone marrow-derived macrophages.

### Metabolomic Profiles of iAs-treated BMDMs

In addition to lipid analytes, we also measured the metabolic profiles of iAs-treated (0, 0.01, or 0.1 µM iAs) BMDMs using LC/MS. Of the 23 metabolites measured (**Supplemental Table 3**), only 3/23 and 6/23 were approaching or at significance in iAs-treated female- and male-derived BMDMs, respectively (**Fig. 8**). At 0.1 µM iAs exposure in females, norepinephrine levels were higher, at near statistical significance (*p*=0.0503). At the highest exposure dose, L-arginine was significantly decreased (*p*<0.01) and S-adenosylmethionine (SAM) was decreased, albeit nonsignificantly (*p*= 0.0933) (**Fig. 8A**). In male BMDMs, significant decreases were observed for L-ascorbic acid (*p*<0.05 at 0.1 µM iAs), norepinephrine (*p*<0.01 for 0.01 µM iAs; *p*<0.05 at 0.1 µM iAs), succinic acid (*p*<0.01 for 0.01 µM iAs; *p*<0.05 at 0.1 µM iAs), L-arginine (*p*<0.01 for 0.01 µM iAs; *p*<0.0001 at 0.1 µM iAs) and γ-Aminobutyric acid (GABA) approaching significance (*p*=0.085 for 0.01 µM iAs) (**Fig. 8B**). These results suggest that iAs treatment suppresses metabolite availability in BMDMs and helps to support the known effects of iAs as a methyl scavenger (as shown by decreases in SAM) and as an activator of Arg1 (as shown by decreases in L-arginine).

**Figure 8.**
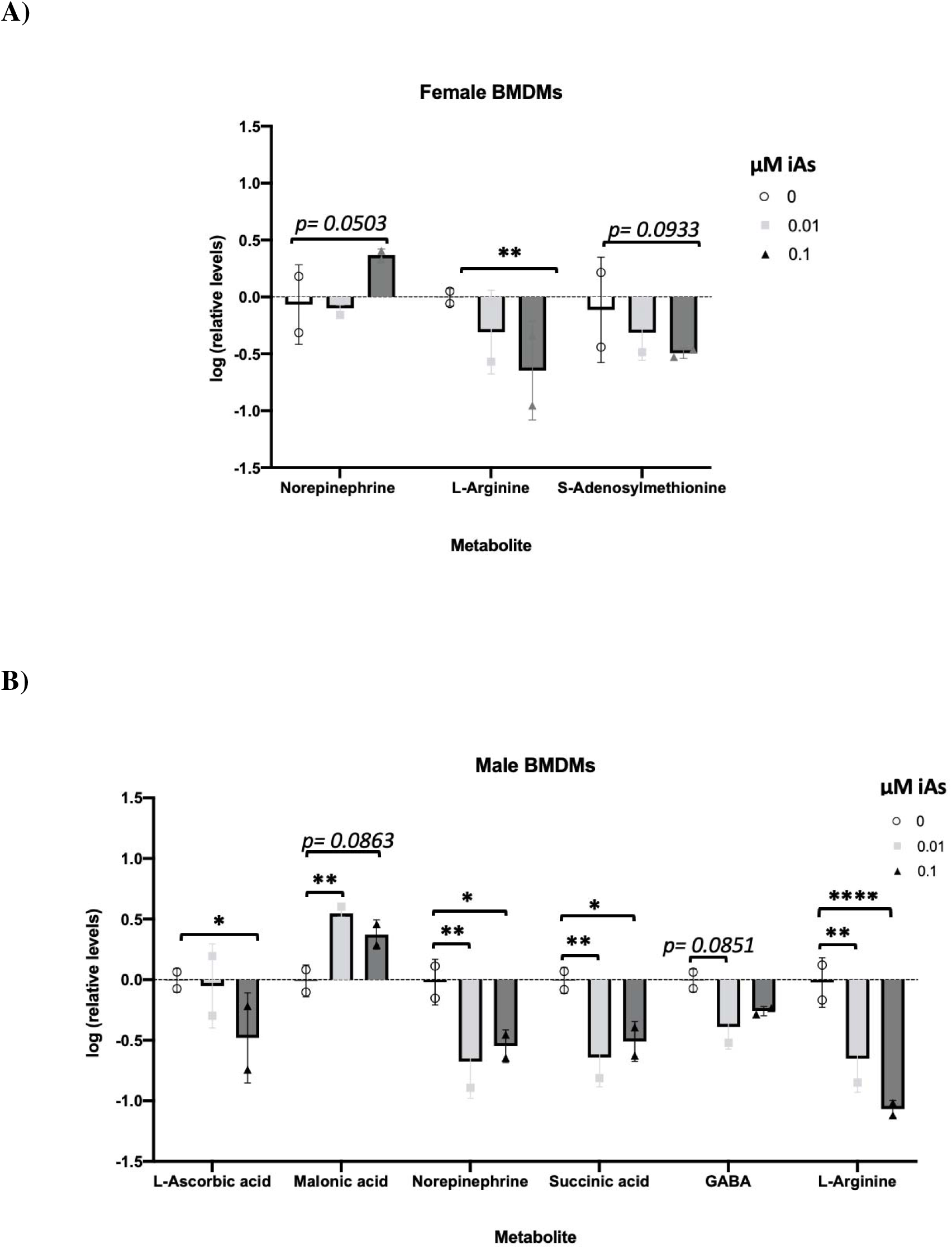
Metabolite profiles of iAs-exposed BMDMs. Polar and nonpolar relative lipid levels from BMDMs exposed to 0, 0.01, or 0.1 µM iAs 1 wk during differentiation were determined by LC/MS. n=2 biological replicates per dose for both males and females, with only significant (or approaching significant) analytes plotted (females: 3/21 analytes vs. males: 6/21 analytes). Plots show mean ± SD. \**p*-value<0.05; \*\**p*-value<0.01; \*\*\**p*-value<0.001; \*\*\*\**p*-value<0.0001. iAs=inorganic arsenic; BMDMs=bone marrow-derived macrophages.

### Migration of iAs-treated Macrophages Toward Cancer Cell-conditioned Media

To contextualize our phenotypic findings within the framework of iAs-associated disease, we next assessed how chronic iAs exposure influenced macrophage responses to cancer cell signals. First, we evaluated how iAs exposure affected macrophages’ ability to migrate toward cancer cell conditioned media. Conditioned media from two murine lung cancer cell lines, LLC1 and KLN205, was used to test the migratory capacity of male and female BMDMs by transwell migration assay. Migration toward LLC1 conditioned media was significantly increased in both male- and female-derived iAs-exposed M0 (unstimulated) BMDMs compared to unexposed M0 controls (*p*<0.01, **Fig. 9A, 9B**). In contrast, migration toward KLN205 conditioned media was not significantly altered in neither male-nor female-derived BMDMs, although iAs-exposed female-derived BMDMs exhibited a noteworthy increase in migration compared to controls (*p*=0.0662, **Fig. 9A**). These results indicate that iAs-treated macrophages migrate more readily toward lung cancer cell-conditioned media compared to controls.

**Figure 9.**
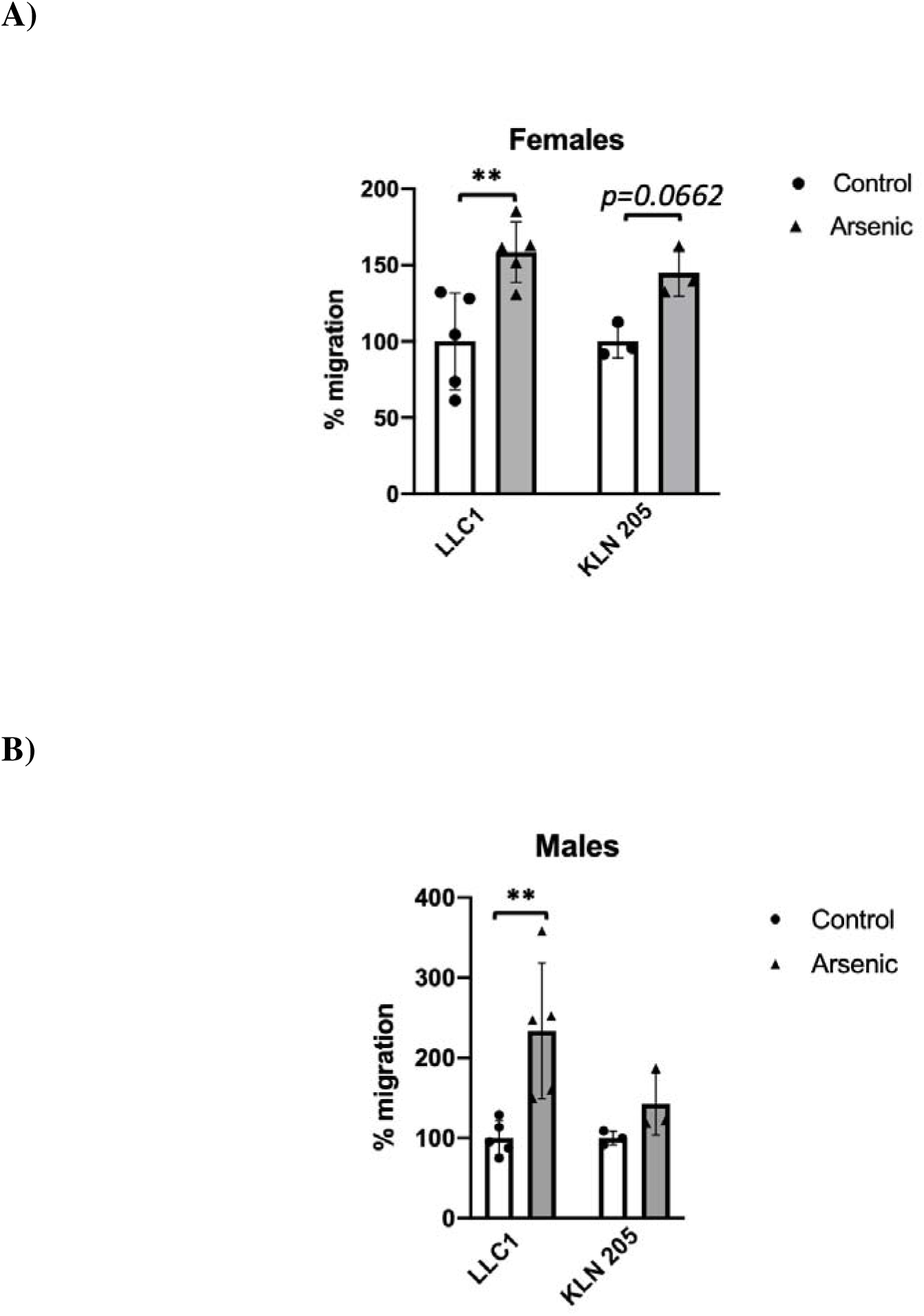
Migration of iAs-exposed BMDMs toward cancer cell conditioned media. Transwell migration assay was used to measure migratory capacity of exposed M0 female and male BMDMs toward LLC1 and KLN205 lung cancer cell conditioned media. n=3-5 biological replicates. Plots show mean ± SD of percent migration measured with crystal violet staining. **p<0.01. iAs= inorganic arsenic; BMDMs= bone marrow-derived macrophages.

### Proliferation of Cancer Cells Cocultured with iAs-treated Macrophages

To test the tumor-promoting potential of iAs-exposed macrophages, we cocultured chronic iAs-exposed RAW 264.7 macrophages with three different cancer cell lines: MLE-15 (lung), LLC1 (lung), or MB49 (bladder). After the coculture period, we determined how the microenvironment created by iAs-exposed RAW 264.7 macrophages drove cancer cell proliferation by MTT assay. MB49 bladder cancer cells cocultured with iAs-exposed RAW 264.7 macrophages exhibited significantly increased proliferation compared to those cocultured with nonexposed macrophages (*p*<0.05, **Fig. 10A**). LLC1 cells also showed increased proliferation in response to coculture with exposed macrophages, though nonsignificant (*p*=0.081, **Fig. 10A**). MLE-15 cells cocultured with iAs-exposed RAW 264.7 macrophages showed no significant proliferation differences compared to those cocultured with nonexposed RAW 264.7 macrophages. However, MLE-15 cells cocultured with female-derived iAs-exposed BMDMs demonstrated a significant increase in proliferation compared to control cocultures (*p*<0.05, **Fig. 10B, 10C).** These results suggest that iAs-treated M0 macrophages create a tumor microenvironment that promotes cancer cell growth.

**Figure 10.**
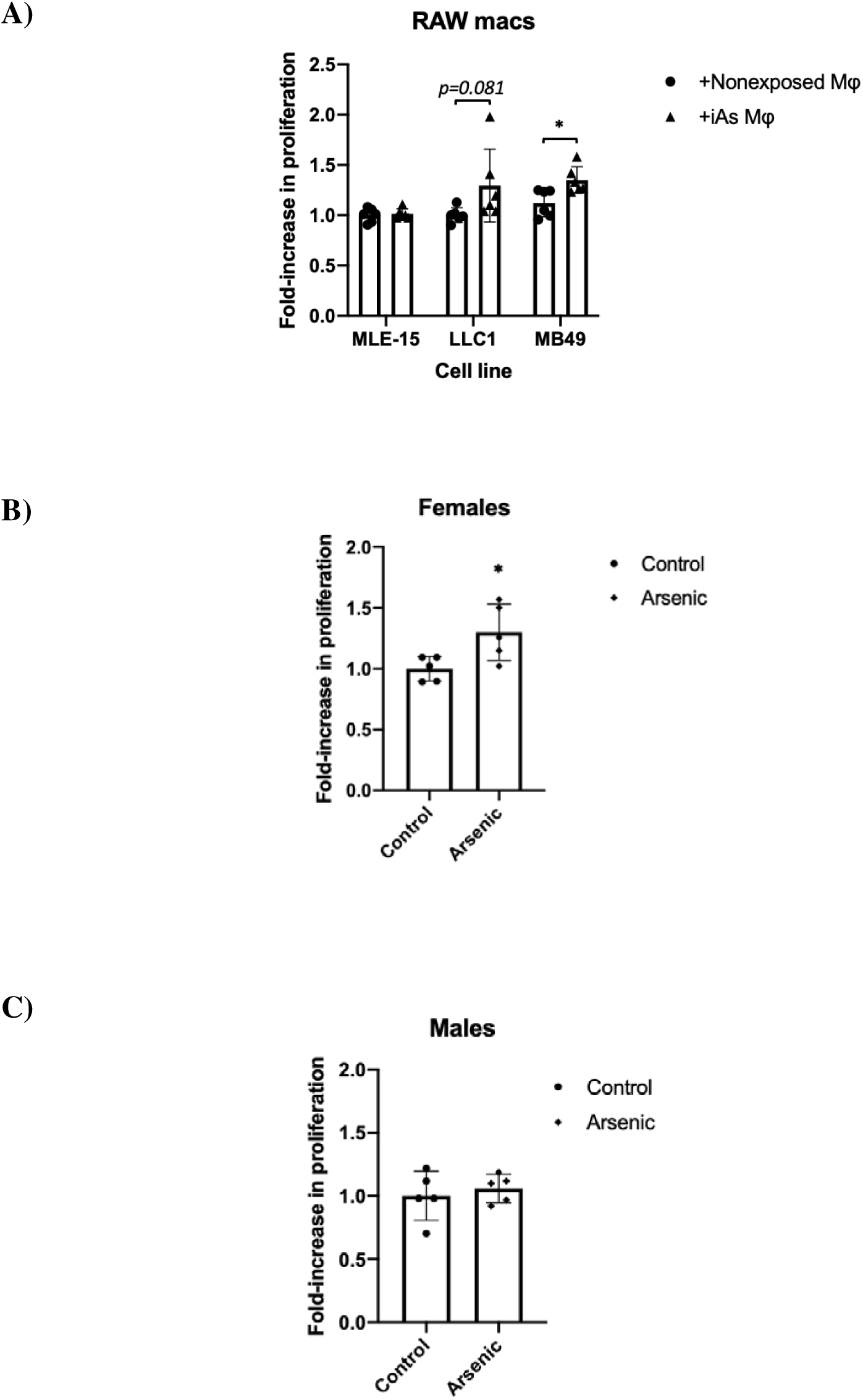
Proliferation of cancer cells cocultured with iAs-exposed macrophages. Proliferation of cancer cells was measured by MTT assay after coculture with exposed M0 RAW 264.7 macrophages (A), female (B) or male (C) BMDMs. n=5 replicates. Plots show mean ± SD of proliferation normalized to control*. *p<0.05.* iAs= inorganic arsenic; BMDMs= bone marrow-derived macrophages.

### Expression of Cancer Stem Cell Markers in Cells Cocultured with iAs-treated RAW 264.7 Macrophages

A subset of replicates of MLE-15, LLC1, and MB49 cancer cells cocultured with iAs-exposed or nonexposed RAW 264.7 macrophages were used for targeted gene expression analysis of cancer stem cell markers. Genes associated with arsenic exposure that were selected for analysis included Yes-Associated-Protein 1, Cyclooxygenase-2 (*YAP* and *SOX2*, respectively) (Michel-Ramirez et al., 2020; Ooki et al., 2018) and self-renewal Notch Receptor 1 (*NOTCH1*) (Tokar et al., 2010). Gap Junction Protein beta 1 (*GJB1)*, a gene involved in cell-cell communication and reported to be downregulated in bladder cancer (Zaravinos et al., 2011) but which has dual roles in other cancer types (Aasen et al., 2019; Yang et al., 2021), was also examined. *NOTCH1* expression was unchanged across all cancer cell lines; however, changes were observed for *YAP*, *SOX2*, and *GJB1*. MB49 bladder cancer cells cocultured with iAs-exposed RAW 264.7 macrophages had significantly upregulated *YAP* expression compared to those cocultured with controls (*p*<0.01, **Fig. 11A**). *SOX2* expression was elevated in both MLE-15 and LLC-1 cells following coculture, but only the increase in MLE-15 cells reached statistical significance (*p*<0.001, **Fig. 11B**; LLC1 *p*=0.123). *GJB1* expression was significantly decreased in LLC1 cells cocultured with iAs-exposed RAW 264.7 macrophages (*p*<0.05, **Fig. 11C**). These results suggest that iAs exposure modulates how M0 macrophages respond to tumor cell conditioned media and that iAs-treated macrophages create a microenvironment supportive of cancer cell growth and cancer stem cell marker expression.

**Figure 11.**
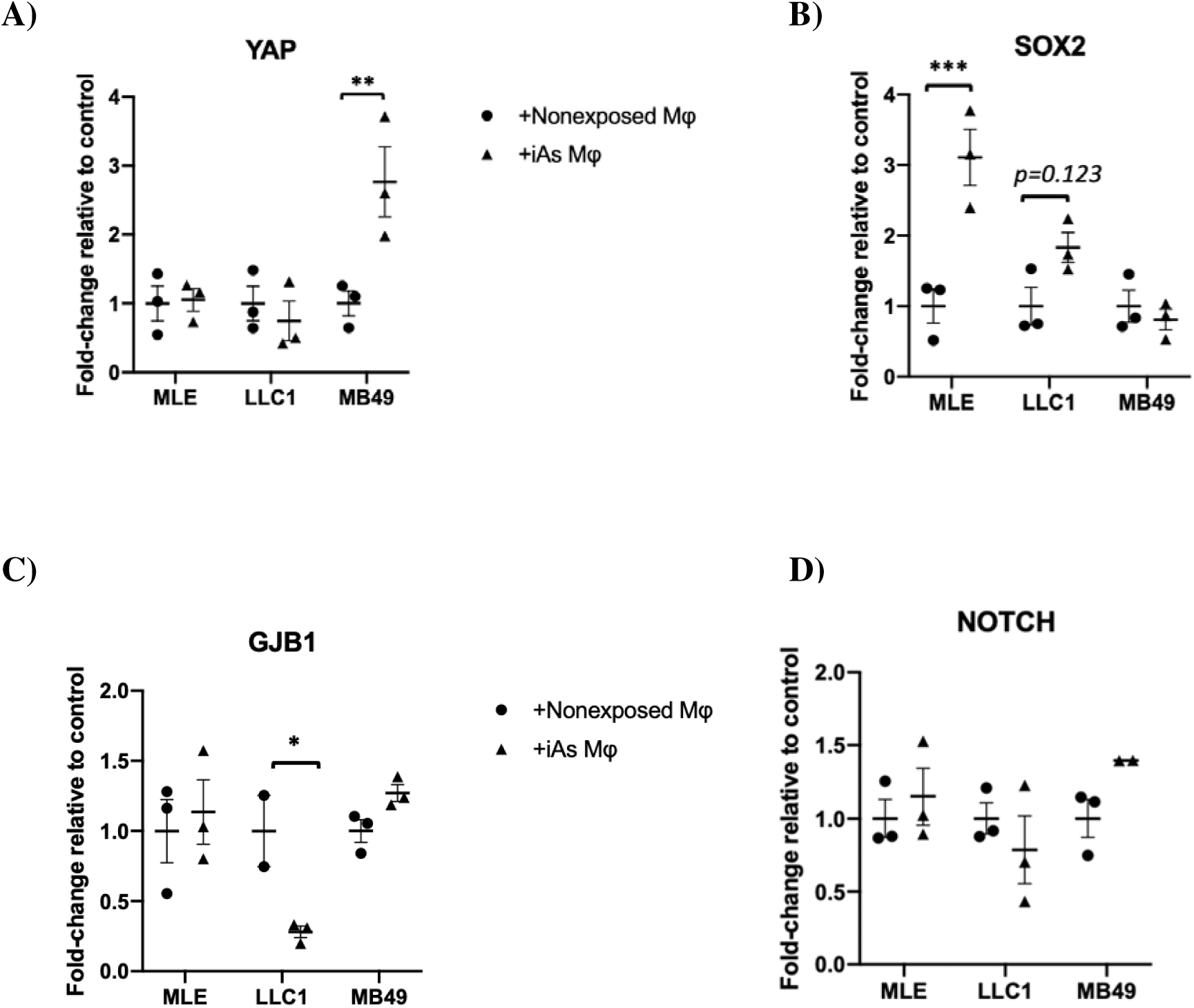
Gene expression analysis of cancer targets in cancer cell lines after coculture with iAs-exposed RAW 264.7 macrophages. RT-qPCR was used to measure expression of select cancer targets YAP (A), SOX2 (B), GJB1 (C), and Notch (D). n=3 replicates. Plots show mean ± SEM of expression normalized to control using GAPDH.

## Discussion

In this study, we report sex- and stimulation-dependent phenotypic changes in macrophages chronically exposed to inorganic arsenic (iAs). The decrease in nitric oxide (NO) production observed in our study is consistent with prior findings from chronic arsenic exposure in humans (Banerjee et al., 2009), as well as murine peritoneal macrophages (Arkusz et al., 2005; Liu et al., 2008), splenic macrophages (Bishayi & Sengupta, 2006; Sengupta & Bishayi, 2002), and RAW 264.7 macrophages (Chakravortty et al., 2001). To our knowledge, however, we are the first to report a decrease in NO production in RAW 264.7 macrophages following chronic exposure to iAs at a dose as low as 0.1 µM (**Fig. 2C**). Given that NO is a major mediator of inflammation and plays an essential role in killing of infectious pathogens and tumor cells (Nath & Kashfi, 2020; Tripathi, 2007), our findings suggest that chronic arsenic exposure dampens the proinflammatory response of macrophages in response to M1 stimuli.

In this study, we demonstrate that, under the conditions of our iAs exposure model, there are significant reductions in the expression of chemokines CCL4, CCL5, and CXCL2. CCL4 and CCL5 were downregulated in M1 and M2 iAs-treated female BMDMs, respectively. These chemokines signal through chemokine receptors (CCRs) CCR1 and CCR5, which are expressed on a wide range of immune cells, including monocytes, macrophages, dendritic cells, T cells, B cells, and natural killer cells (Mantovani et al., 2004), and function to increase immune cell adhesion to endothelial cells and elicit a pro-inflammatory response (Gunawardana et al., 2021; Quandt & Dorovini-Zis, 2004). Similarly, CXCL2 signals via CXCR2 on neutrophils to trigger adhesion and migration through the vascular endothelium (Wang et al., 2021), and its reduction may compromise the ability to mount a pro-inflammatory immune response.

Decreases in TIMP-1 and CHI3L1 were observed in iAs-treated M0 female and male macrophages, respectively. TIMP-1 is an endogenous inhibitor of matrix metalloproteinases (Winkler et al., 2020) which may be relevant in iAs-induced diseases like cancer, where macrophages act as major remodelers of the extracellular matrix to promote angiogenesis and modulate T cell infiltration (Afik et al., 2016; Gunawardana et al., 2021). CHI3L1 (chitinase 3 like 1) is a member of the glycosyl hydrolase 18 family that lacks chitinase activity. CHI3L1 is expressed in both M1 and M2 macrophages depending on stimulation context (Zhao et al., 2020) and signals to regulate growth and differentiation of macrophages and other cell types. Like TIMP-1, there is also evidence that CHI3L1 regulates remodeling of the extracellular matrix (Kzhyshkowska et al., 2006; Li et al., 2021). These findings suggest that arsenic exposure may impair macrophages’ ability to remodel the extracellular matrix. Future experiments could explore this *in vitro* with biological scaffold techniques (Valentin et al., 2009) and *in vivo* in the context of disease with histological analysis targeting markers involved in remodeling (Rahman et al., 2020).

Because arsenic can also disrupt cell differentiation (Lemarie et al., 2006; Liu et al., 2019; Sakurai et al., 2006), we also wanted to characterize MDSCs that were derived from our bone marrow cultures. MDSCs are potent immunoregulators and precursors of macrophages that are also elicited from bone marrow. They have implicated roles in arsenic-relevant diseases, including cancers (Yang et al., 2020) and atherosclerosis (Foks et al., 2016). However, to our knowledge, no direct link between arsenic exposure and changes in MDSC phenotype or function has been reported. Unique to our MDSCs, we observed a decrease of PD-L1 expression on M1 MDSCs and an increase in cells expressing PD-L1 in our M2-stimulated MDSCs (**Supp. Figs. 1 and 2**). PD-L1 is a common marker of an immunosuppressive microenvironment in cancer and is expressed by macrophages to dampen the inflammatory response and inhibit T cell effector function (Jiang et al., 2019; Liu et al., 2020). These findings suggest that chronic iAs exposure reprograms not only macrophages but also their myeloid precursors toward a less inflammatory, more immunosuppressive phenotype.

Intracellular lipid droplets provide sources of fatty acids for macrophages, and fatty acid metabolism is tightly linked to metabolic rewiring of M1 and M2 macrophages. For example, M1 macrophages have greater metabolic flux through aerobic glycolysis, the pentose phosphate pathway, and increased fatty acid synthesis to fuel the energetic demand of M1-associated functions such as phagocytosis and rapid generation of inflammatory mediators (Batista-Gonzalez et al., 2019). M2 macrophages, on the other hand, rely heavily on glutaminolysis, the TCA cycle, oxidative phosphorylation (OXPHOS), and fatty acid oxidation (Ren et al., 2019; Sadiku & Walmsley, 2019). We first observed an increase in lipid droplet formation in chronic iAs-esposed RAW 264.7 macrophages using a fluorescent lipid staining assay (**Fig. 6C, 6D**).

To identify which specific lipids were upregulated in iAs-exposed macrophages, we performed LC/MS lipidomics analysis on BMDMs treated with MMA(III), a toxic metabolite of iAs. We identified several lipid classes that were upregulated, including prostaglandins, ether lipids, lysophospholipids, and sphingolipids (**Fig. 7**). These findings suggest that arsenic exposure disrupts the arachidonic acid metabolism cascade, with potential implications for macrophage function in disease. For example, prostaglandins are derived from arachidonic acid through enzymatic activity of COX2 and signal in an autoregulatory fashion to downregulate proinflammatory (e.g., TNF) cytokine production and upregulate anti-inflammatory (e.g., IL-10) cytokine production in macrophages (Saleh et al., 2020). Aditionally, prostaglandins participate bi-directionally in cancer cell-macrophage crosstalk as prostaglandins secreted from macrophages can also serve as ligands to G-protein coupled receptors (GPCRs) on cancer cells to promote survival, migration, and further inflammation and immunosuppression (Inaba et al., 2003; Kashiwagi et al., 2018; Thompson, 2020). Phospholipids have additionally been shown to regulate macrophage phenotype. This has been characterized especially in obesity (Morgan et al., 2021; Poledne et al., 2019; Serbulea et al., 2018) and in cardiovascular diseases like atherosclerosis (Nishikawa et al., 2015; Rosenblat et al., 2006). Macrophage-derived phospholipids are also upregulated in macrophage-cancer cell coculture systems (Rabold et al., 2020). We observed elevated sphingosine-1-phosphate (S1P), which is tightly linked to eicosanoid metabolism and has been shown to promote differentiation of monocytes to macrophages and tumor cell signaling (Weigert et al., 2019). However, increased S1P (**Fig. 7**) does not does not coincide with any significantly increased monocyte-macrophage differentiation based on our flow cytometric analysis (**Fig. 4D**). Importantly, S1P may produce paracrine signaling to other cells to regulate migration, proliferation, and differentiation (Weigert et al., 2019).

Interestingly, COX2, a major regulator of arachidonic acid metabolism and the rate-limiting step in prostagladin synthesis, is regulated by S-nitrosylation and may interact directly with iNOS (Salvemini et al., 2013). Production of metabolites downstream of COX2 can suppress NO and NOS isoform production through negative feedback mechansims (Cheng et al., 2002; Mollace et al., 2005). In line with this, we observed reduced NO in our chronic iAs-exposed RAW 264.7 macrophages (**Fig. 2**), and decreased iNOS in our flow cytometric analysis of iAs-exposed BMDMs (**Fig. 5E**). This connection represents one plausible explanation for these collective phenotypic findings.

We also observed distinct effects on iAs-exposed male-derived BMDMs, which had significant reductions in L-ascorbic acid, norepinephrine, and succinic acid (**Fig. 8B**). While succinic acid is associated with M2 polarization and flux through the TCA cycle (Trauelsen et al., 2021), ascorbate has dual effects on macrophage polarization and function; it has been shown to attenuate LPS-induced activation and cytokine production in monocytes (Schmidt et al., 2022), partially explaining the iAs-induced cytokine and chemokine profiles we observed (**Fig. 3**). Nevertheless, it also diminishes expression of certain M2 markers including CD163, which contradicts our flow cytometric analysis (**Fig. 5B**), further supporting the mixed polarization phenotype.

In addition to determining migratory capability of iAs-exposed macrophages toward lung cancer cell conditioned media, we cocultured either chronic exposed RAW 264.7 macrophages in a transwell system (without cell-cell contact) with MLE-15, LLC1, or MB49 cells to further understand arsenic’s effects on macrophages in the context of cancer promotion. Interestingly, both LLC1 lung cancer cells and MB49 bladder cancer cells displayed increased proliferation after coculture with iAs-exposed macrophages compared to controls, with MB49 proliferation reaching statistical significance (**Fig. 10A**). Of note, while we have not tested their ability to promote proliferation of other cancer cell types, iAs-exposed female BMDMs cocultured with the tumorigenic lung epithelial cell line MLE-15 promoted their proliferation in a statistically significant manner (**Fig. 10B**). Because this coculture does not allow for cell-cell contact, we can infer that the presence of signaling molecules derived from iAs-exposed macrophages in the shared transwell media drove the proliferation of cancer cells. This is not surprising, given that macrophage-derived soluble mediators such as metabolites, chemokines, and cytokines (Ge & Ding, 2020), as well as non-soluble mediators such as lipids and the content of exosomes (Liu et al., 2020), have been shown to influence tumor cell growth. We next sought to determine which cancer-related genes might be driving the increased proliferation after coculture with arsenic-exposed macrophages. We harvested cancer cells after coculture and performed gene expression analysis for a panel of PCR primers (**Supplemental Table 5**). Subsequent qPCR analysis revealed cancer stem cell markers *YAP* and *SOX2* significantly increased in MB49 and MLE-15 cells, respectively, after coculture with iAs-exposed macrophages compared to controls (**Fig. 11A, 11B**). Together, these results imply that iAs-exposed macrophages create a microenvironment conducive to cancer cell growth and cancer stem cell promotion. Indeed, many of the properties we have identified in our analysis of iAs-exposed macrophages have been characterized in TAMs, including expression of M2 markers (**Figs. 4** and **5**) as negative prognostic factors, as well as enhanced lipid biosynthesis (Rabold et al., 2020) (**Figs. 6** and **7**).

Our study has several strengths. First, our exposure model uses a human-relevant low-dose arsenic exposure level. Even though we show that this dose is noncytotoxic to M0 BMDMs (**Fig. 1**), we also normalize to cell viability and cell number whenever possible to minimize off-target results from dead or dying cells because macrophages are also potent responders to Damage-associated molecular patterns (DAMPs) (Blander, 2017) and plating density alone can influence their differentiation states (Lee & Hu, 2013). We also conducted sex stratified analyses when using BMDMs since there are differences between female and male biotransformation of iAs (Lindberg et al., 2008; Muhetaer et al., 2022; Pierce et al., 2013) and sex differences in morbidity and mortality of iAs-induced diseases, especially cancers (Ferreccio et al., 2000; Parvez et al., 2013; M. Rahman et al., 2006; Smith et al., 2006). In addition, macrophage phenotype is influenced by sex through mechanisms such as hormone signaling (Barcena et al., 2021; Klein & Flanagan, 2016) and sex-linked genes (Schurz et al., 2019; Syrett et al., 2019). Combined with these sex differences reported in the literature, the sex-dependent changes in iAs-exposed macrophages that we observed in this study across cytokine profiles (**Fig. 3**), expression of polarization markers (**Figs. 4** and **5**), and influence on cancer cell growth (**Fig. 10**) support the need for further investigation into sex-informed toxicology. Another strength of our study design is that when characterizing our arsenic-exposed macrophages, we utilized M0, M1, and M2 stimulation conditions for multiple endpoints. This allowed us to contextualize changes in NO production, chemokine and cytokine production, and flow cytometric characterization of polarization markers, as many of these changes are stimulation-dependent (**Figs. 2–5**). Still, we recognize that this contextualization is limited compared to the mixed stimulation conditions BMDMs experience *in vivo.* In the future, experiments should include cancer cell conditioned media as its own stimulation control.

One limitation of this study is that thus far, we have only explored the tumor-promoting effects of shared conditioned media on cancer cell promotion. Future work will continue to pursue the mechanism by which exposed macrophages drive cancer cell proliferation with targeted agonists and inhibitors based on our phenotypic findings and knowledge of upstream ligands that signal to regulate cancer stem cell marker expression. For example, we will target enzymes involved in lipid synthesis (specifically to upregulated targets we observed in our lipidomics analysis), such as COX2, a key enzyme in prostaglandin synthesis (e.g., with inhibitor Celecoxib) (Zhang et al., 2016), phospholipase A2, a key enzyme regulating phospholipid production (e.g., with inhibitor Varespladib) (Arsenault et al., 2011) and S1P sphingosine kinase, a key enzyme regulating S1P production (e.g., with inhibitor SKI-II) (Pitman et al., 2022).

Another limitation of this study is that we were only to run lipidomics/metabolomics analyses on macrophages under a single stimulation condition (M0 female BMDMs) with MMA(III) exposure rather than with exposure to sodium (meta) arsenite, which was used in our exposure model for other functional endpoints. Since macrophages have distinct modes of lipid metabolism upon receiving polarization queues and in many disease statuses (Batista-Gonzalez et al., 2019; Kelly & O’Neill, 2015), we need to further explore the effects of arsenic under additional M1 and M2 polarization conditions across sexes.

In summary, iAs does not drive unidirectional polarization shift but induces a hybrid macrophage phenotype with sex dependent metabolic, phenotypic, and functional changes. These macrophages contribute to tumor-promoting microenvironment and represent a key immunotoxicological target in arsenic-associated diseases. Overall, understanding the relationship between iAs-relevant diseases and macrophage biology may offer insight toward novel strategies for macrophage-associated disease therapy.

## SUPPLEMENTAL FIGURES

**Supplemental Figure 1.**
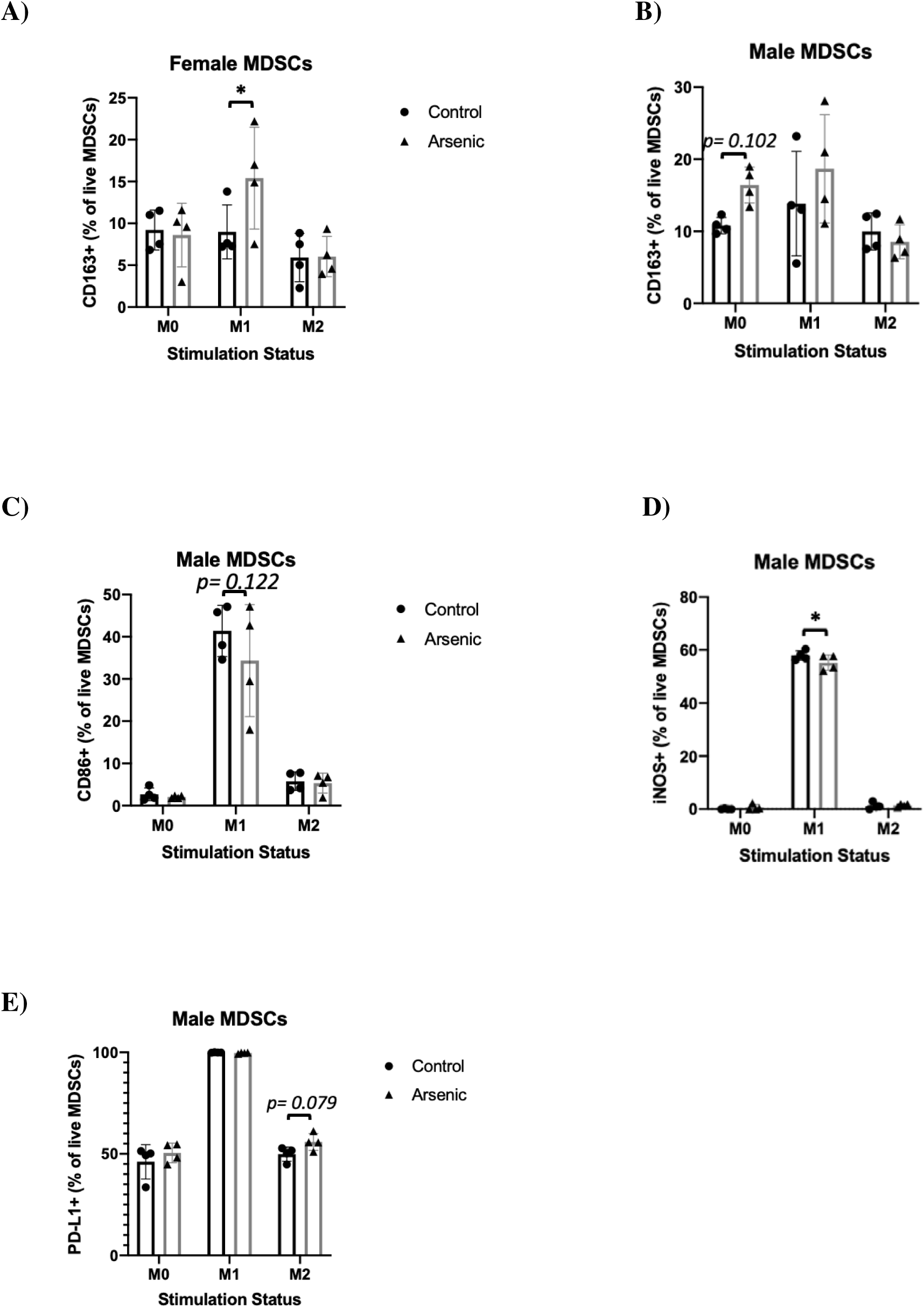
Flow cytometric characterization of iAs-exposed female and male MDSCs as a frequency of parent gates. MDSCs (Ly6C+CD11b+F4/80-MHCII-live cells) were analyzed from BMDM cultures for expression of select polarization and functional markers CD163 (A B), CD86 (C), iNOS (D), and PD-L1 (E). Only plots with *p*-values <0.15 are included (complete panel with all markers analyzed can be found in Supplemental Table 4). Plots show % expression of parent gate on the y-axis by stimulation condition on the x-axis, with the mean ± SD. n=4 biological replicates. \**p*-value<0.05. iAs=inorganic arsenic; MDSCs=monocytic myeloid derived suppressor cells.

**Supplemental Figure 2.**
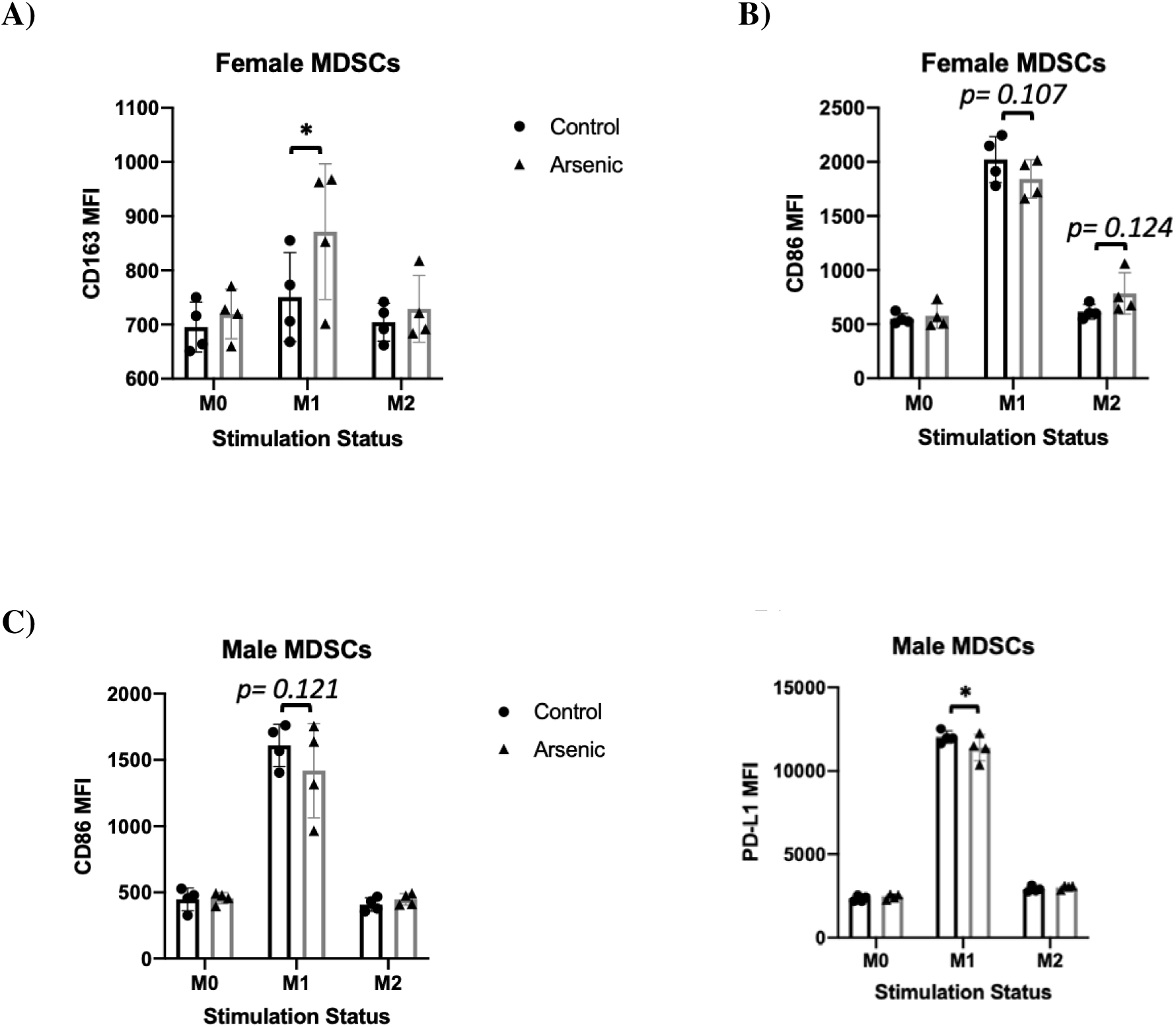
Flow cytometric characterization of iAs-exposed female and male MDSC polarization and functional markers as MFI expression levels. MDSCs (Ly6C+CD11b+F4/80-MHCII-live cells) were analyzed from BMDM cultures for expression of select polarization and functional markers CD163 (A), CD86 (B, C), and PD-L1 (D). Only plots with *p*-values <0.15 are included (complete panel with all markers analyzed can be found in Supplemental Table 4). Plots show MFI on the y-axis by stimulation condition on the x-axis, with the mean ± SD. n=4 biological replicates. \**p*-value<0.05; \*\**p*-value<0.01. iAs=inorganic arsenic; MDSCs=monocytic myeloid derived suppressor cells. MFI= mean fluorescence intensity.

**Supplemental Figure 3.**
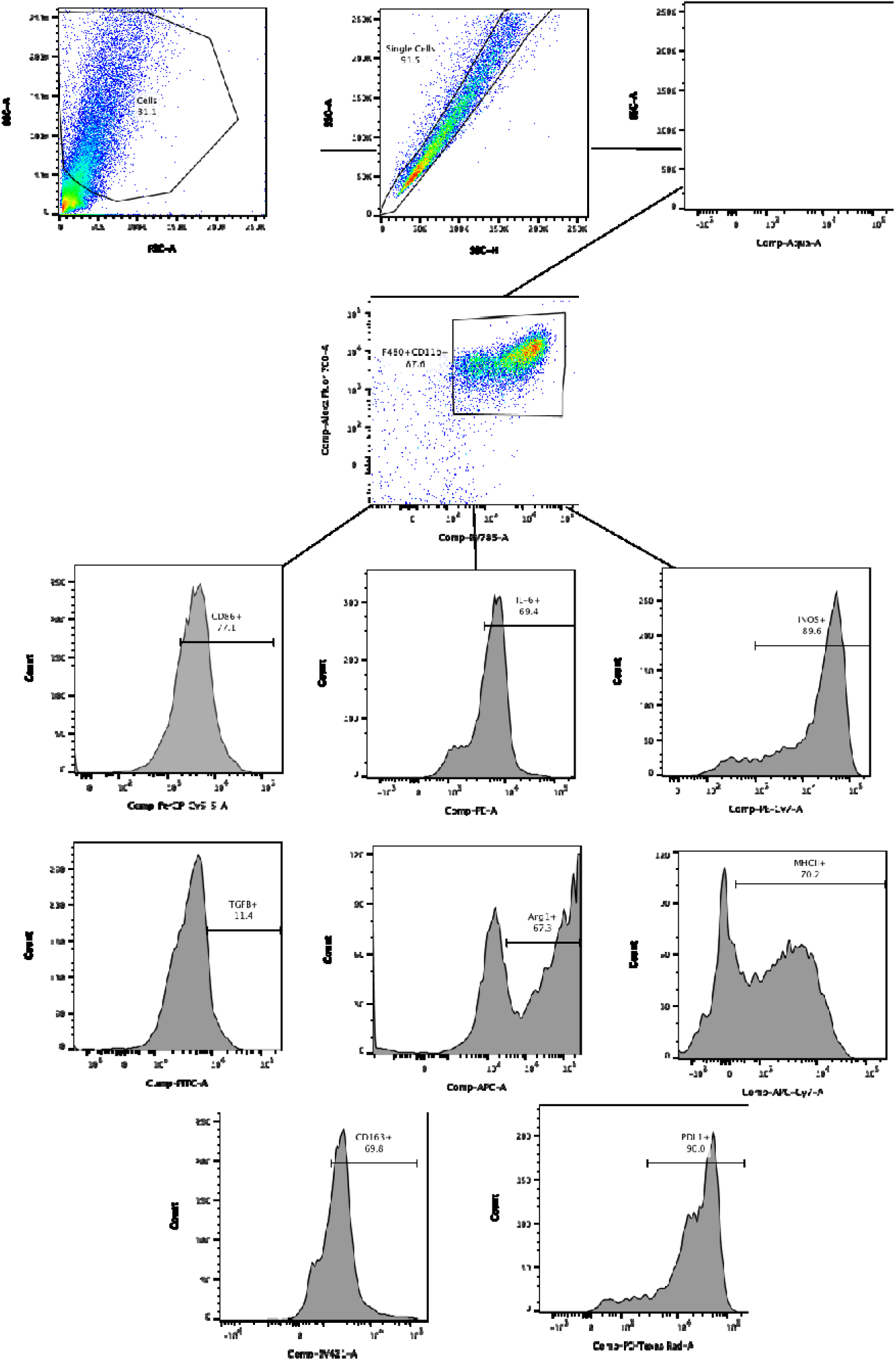
Representative flow cytometry gating strategy.

## SUPPLEMENTAL TABLES

**Supplemental Table 1.** Cytokine targets.

| Target Name |
| --- |
| BAFF/BLyS/TNFSF13B |
| CCL11/Eotaxin |
| CCL19/MIP-3 beta |
| CCL2/JE/MCP-1 |
| CCL3/MIP-1alpha |
| CCL4/MIP-1 beta |
| CCL5/RANTES |
| CCL7/MCP-3/MARC |
| CXCL1/GRO alpha/KC/CINC-1 |
| Chitinase 3-like 1 |
| EGF |
| FGF basic/FGF2/bFGF |
| GDF-15 |
| GM-CSF |
| IFN-gamma |
| IL-1 alpha/IL-1F1 |
| IL-10 |
| IL-13 |
| IL-16 |
| IL-17/IL-17A |
| IL-2 |
| IL-27 |
| IL-3 |
| IL-4 |
| IL-6 |
| IL-7 |
| LDLR |
| LIX |
| TIMP-1 |
| VEGF |
| CXCL2/MIP-2 |
| G-CSF |
| IL-12p70 |
| IL-1b |
| IL-5 |
| TNF |

**Supplemental Table 2.** Lipid targets.

| Lipid class | Target Name |
| --- | --- |
| Acyl carnitines | C:12 (lauroyl carnitine)<br>C16:0 (palmitoyl L-carnitine) |
| Eicosanoids | PGE<sub>2</sub> (Prostaglandin E<sub>2</sub>) |
| Ether lipids | C16:0 (lyso PAF)<br>C18:0 (lyso PAF)<br>C18:1 (lyso PAF)<br>C16:0/20:4 (phosphatidic acid ether)<br>C16:0 LPCe (phosphocholine ether)<br>C18:1 LPCe (phosphocholine ether)<br>C18:0/C18:1 (phosphotidylglycerol ether) |
| Lysophospholipids | C20:4 LPA (lysophosphatidic acid)<br>C20:4 LPC (lysophosphatidylcholine)<br>C18:1 LPE (lysophosphoethanolamine)<br>C18:0 LPE (lysophosphoethanolamine)<br>C20:4 LPE (lysophosphoethanolamine)<br>C18:0 LPS (lysophosphoethanolamine) |
| Phospholipids | C16:0/C18:1 PA<br>C16:/C20:4 PA<br>C18:0/C18:1 PA<br>C18:0/C20:4 PA<br>C16:0/C20:4 PC<br>C18:0/C18:1PC<br>C18:0/C20:4 PC<br>C16:0/C20:4 PE<br>C18:0/C20:4 PE<br>C16:0/C20:4 PS |
| Sphingolipids | C16:0 CM<br>C18:0 CM<br>C20:4 SM<br>C16:0 S1P |

**Supplemental Table 3.** Metabolite targets.

| Target Name |
| --- |
| D-(-)-β-Hydroxybutyric acid |
| D-α-Hydroxyglutaric acid |
| L-Ascorbic acid |
| Malonic acid |
| N-Acetylaspartate |
| Norepinephrine (noradrenaline) |
| Succinic acid |
| Uric acid |
| Xanthosine |
| 5-HIAA / 5-Hydroxy-3-indoleacetic acid |
| Acetylcholine |
| Choline |
| Creatine |
| GABA / gamma-Aminobutyric acid |
| GSH / Glutathione |
| GSSG / L-Gluthathione (oxidized) |
| L-Arginine |
| L-Glutamic acid |
| L-Histidine |
| L-Methionine |
| L-Tryptophan |
| S-Adenosylmethionine |
| SAH / S-Adenosyl-L-homocysteine |

**Supplemental Table 4.** Flow cytometric panel design.

| Flow cytometric panel design for macrophages and MDSCs |  |  |  |
| --- | --- | --- | --- |
| Antibody | Clone | Conjugate | Company |
| Viability | - | Aqua | Biolegend |
| F4/80 | BM8 | BV785 | Biolegend |
| Ly6C | HK1.4 | BV605 | Biolegend |
| CD11b | M1/70 | AF700 | Biolegend |
| CD86 | GL-1 | PerCP/Cyanine5.5 | Biolegend |
| CD163 | TNKUPJ | SB436 | eBioscience |
| I-A/I-E (murine MHC II) | M5/114.15.2 | APC/Fire750 | Biolegend |
| PD-L1 (CD274) | 10F.9G2 | PE/Dazzle™ 594 | Biolegend |
| iNOS | CXNFT | PE-Cy7 | eBioscience |
| Arg1 | A1exF5 | APC | eBioscience |
| TGF-β1 (LAP) | TW7-16B4 | FITC | Biolegend |
| IL-6 | MP5-20F3 | PE | Biolegend |

**Supplemental Table 5.**
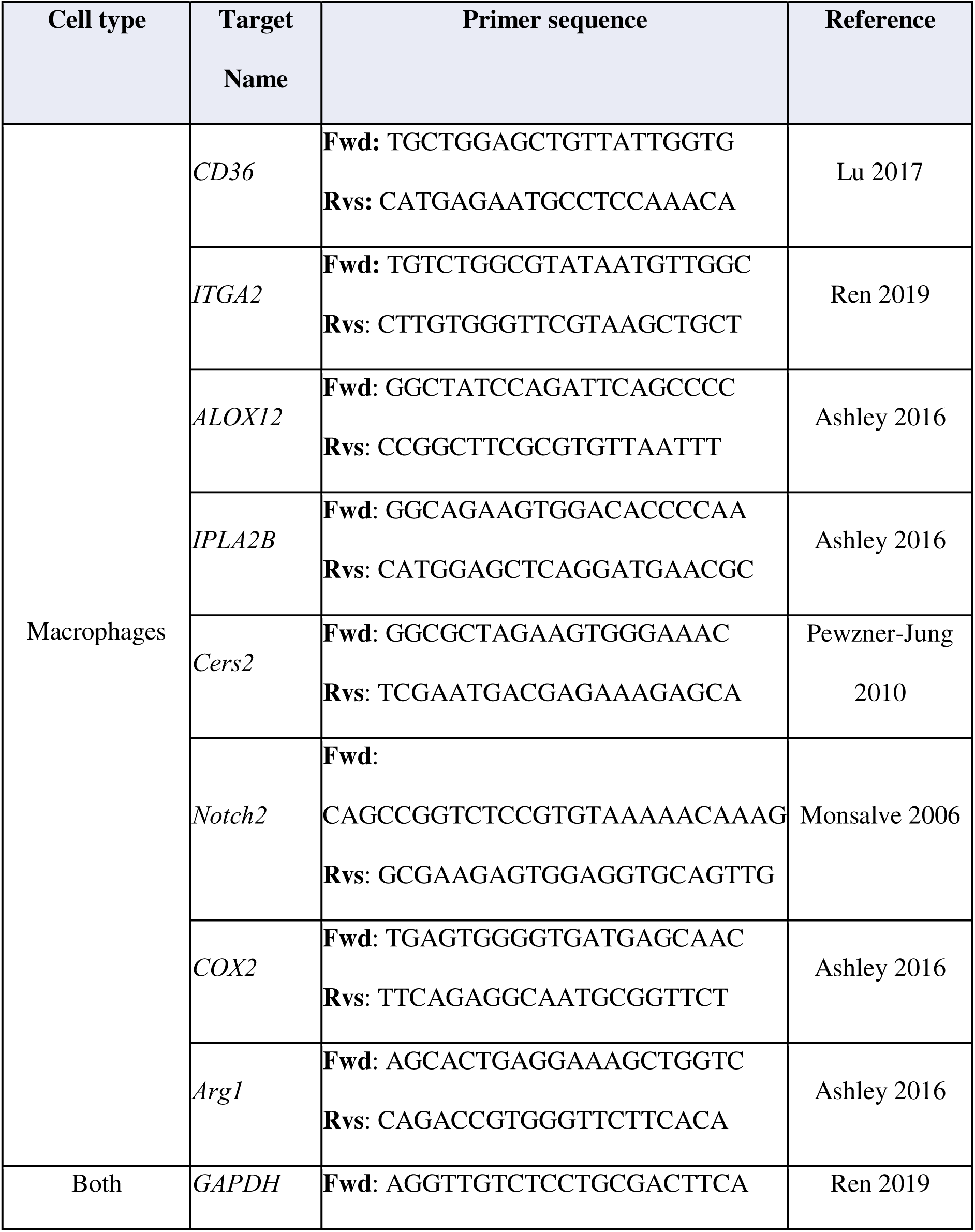
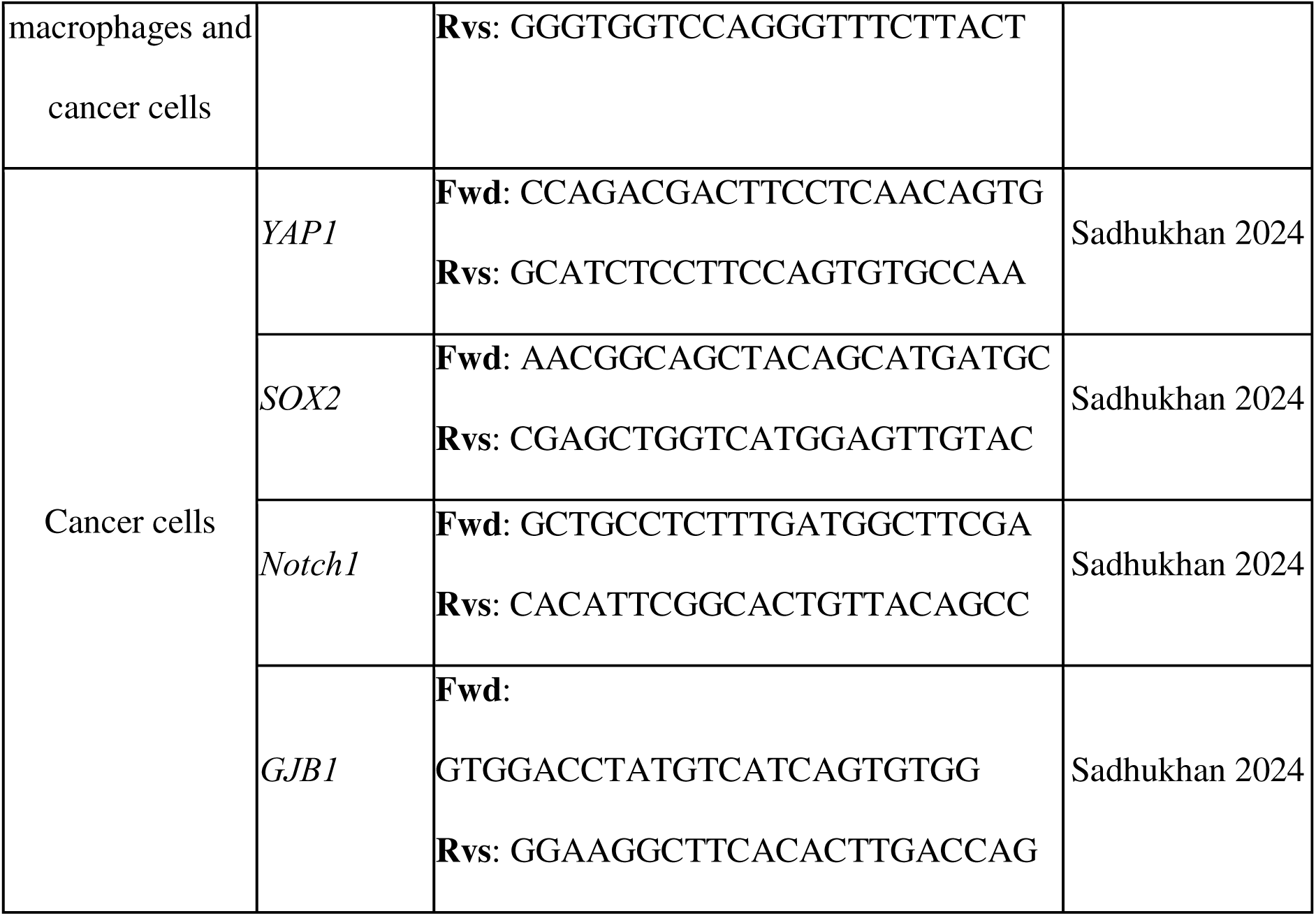
Primers for RT-qPCR validation.

## References

Aasen, T., Sansano, I., Montero, M. Á., Romagosa, C., Temprana-Salvador, J., Martínez-Marti, A., Moliné, T., Hernández-Losa, J., & Ramón y Cajal, S. (2019). Insight into the Role and Regulation of Gap Junction Genes in Lung Cancer and Identification of Nuclear Cx43 as a Putative Biomarker of Poor Prognosis. Cancers, 11(3), 320. 10.3390/cancers11030320

Afik, R., Zigmond, E., Vugman, M., Klepfish, M., Shimshoni, E., Pasmanik-Chor, M., Shenoy, A., Bassat, E., Halpern, Z., Geiger, T., Sagi, I., & Varol, C. (2016). Tumor macrophages are pivotal constructors of tumor collagenous matrix. The Journal of Experimental Medicine, 213(11), 2315–2331. 10.1084/jem.20151193

Amano, F., & Noda, T. (1995). Improved detection of nitric oxide radical (NO.) production in an activated macrophage culture with a radical scavenger, carboxy PTIO and Griess reagent. FEBS Letters, 368(3), 425–428. 10.1016/0014-5793(95)00700-j

Arkusz, J., Stańczyk, M., Lewińiska, D., & Stepnik, M. (2005). Modulation of murine peritoneal macrophage function by chronic exposure to arsenate in drinking water. Immunopharmacology and Immunotoxicology, 27(2), 315–330. 10.1081/iph-200067947

Arsenault, B. J., Boekholdt, S. M., & Kastelein, J. J. P. (2011). Varespladib: Targeting the inflammatory face of atherosclerosis. European Heart Journal, 32(8), 923–926. 10.1093/eurheartj/ehq385

Assouvie, A., Daley-Bauer, L. P., & Rousselet, G. (2018). Growing Murine Bone Marrow-Derived Macrophages. Methods in Molecular Biology (Clifton, N.J.), 1784, 29–33. 10.1007/978-1-4939-7837-3_3

Attreed, S. E., Navas-Acien, A., & Heaney, C. D. (2017). Arsenic and Immune Response to Infection During Pregnancy and Early Life. Current Environmental Health Reports, 4(2), 229–243. 10.1007/s40572-017-0141-4

Bahari, A., & Salmani, V. (2017). Environmentally relevant dose of arsenic interferes in functions of human monocytes derived dendritic cells. Toxicology Letters, 275, 118–122. 10.1016/j.toxlet.2017.05.005

Banerjee, N., Banerjee, S., Sen, R., Bandyopadhyay, A., Sarma, N., Majumder, P., Das, J. K., Chatterjee, M., Kabir, S. N., & Giri, A. K. (2009). Chronic arsenic exposure impairs macrophage functions in the exposed individuals. Journal of Clinical Immunology, 29(5), 582–594. 10.1007/s10875-009-9304-x

Barcena, M. L., Niehues, M. H., Christiansen, C., Estepa, M., Haritonow, N., Sadighi, A. H., Müller-Werdan, U., Ladilov, Y., & Regitz-Zagrosek, V. (2021). Male Macrophages and Fibroblasts from C57/BL6J Mice Are More Susceptible to Inflammatory Stimuli. Frontiers in Immunology, 12, 758767. 10.3389/fimmu.2021.758767

Batista-Gonzalez, A., Vidal, R., Criollo, A., & Carreño, L. J. (2019). New Insights on the Role of Lipid Metabolism in the Metabolic Reprogramming of Macrophages. Frontiers in Immunology, 10, 2993. 10.3389/fimmu.2019.02993

Becker, S. M., & McCoy, K. L. (2003). Gallium arsenide selectively up-regulates inflammatory cytokine expression at exposure site. The Journal of Pharmacology and Experimental Therapeutics, 307(3), 1045–1053. 10.1124/jpet.103.057919

Bellamri, N., Morzadec, C., Fardel, O., & Vernhet, L. (2018). Arsenic and the immune system. Current Opinion in Toxicology, 10, 60–68. 10.1016/j.cotox.2018.01.003

Benjamin, D. I., Cozzo, A., Ji, X., Roberts, L. S., Louie, S. M., Mulvihill, M. M., Luo, K., & Nomura, D. K. (2013). Ether lipid generating enzyme AGPS alters the balance of structural and signaling lipids to fuel cancer pathogenicity. Proceedings of the National Academy of Sciences of the United States of America, 110(37), 14912– 14917. 10.1073/pnas.1310894110

Berghaus, L. J., Moore, J. N., Hurley, D. J., Vandenplas, M. L., Fortes, B. P., Wolfert, M. A., & Boons, G.-J. (2010). Innate immune responses of primary murine macrophage-lineage cells and RAW 264.7 cells to ligands of Toll-like receptors 2, 3, and 4. Comparative Immunology, Microbiology and Infectious Diseases, 33(5), 443–454. 10.1016/j.cimid.2009.07.001

Bishayi, B., & Sengupta, M. (2003). Intracellular survival of Staphylococcus aureus due to alteration of cellular activity in arsenic and lead intoxicated mature Swiss albino mice. Toxicology, 184(1), 31–39. 10.1016/s0300-483x(02)00549-8

Bishayi, B., & Sengupta, M. (2006). Synergism in immunotoxicological effects due to repeated combined administration of arsenic and lead in mice. International Immunopharmacology, 6(3), 454–464. 10.1016/j.intimp.2005.09.011

Blander, J. M. (2017). The many ways tissue phagocytes respond to dying cells. Immunological Reviews, 277(1), 158–173. 10.1111/imr.12537

Bronte, V., Brandau, S., Chen, S.-H., Colombo, M. P., Frey, A. B., Greten, T. F., Mandruzzato, S., Murray, P. J., Ochoa, A., Ostrand-Rosenberg, S., Rodriguez, P. C., Sica, A., Umansky, V., Vonderheide, R. H., & Gabrilovich, D. I. (2016). Recommendations for myeloid-derived suppressor cell nomenclature and characterization standards. Nature Communications, 7(1), 12150. 10.1038/ncomms12150

Carter, D. E., Aposhian, H. V., & Gandolfi, A. J. (2003). The metabolism of inorganic arsenic oxides, gallium arsenide, and arsine: A toxicochemical review. Toxicology and Applied Pharmacology, 193(3), 309–334. 10.1016/j.taap.2003.07.009

Chakravortty, D., Kato, Y., Sugiyama, T., Koide, N., Mu, M. M., Yoshida, T., & Yokochi, T. (2001). The inhibitory action of sodium arsenite on lipopolysaccharide-induced nitric oxide production in RAW 267.4 macrophage cells: A role of Raf-1 in lipopolysaccharide signaling. Journal of Immunology (Baltimore, Md.: 1950), 166(3), 2011–2017. 10.4049/jimmunol.166.3.2011

Chen, F., Chen, J., Yang, L., Liu, J., Zhang, X., Zhang, Y., Tu, Q., Yin, D., Lin, D., Wong, P.-P., Huang, D., Xing, Y., Zhao, J., Li, M., Liu, Q., Su, F., Su, S., & Song, E. (2019). Extracellular vesicle-packaged HIF-1α-stabilizing lncRNA from tumour-associated macrophages regulates aerobic glycolysis of breast cancer cells. Nature Cell Biology, 21(4), 498–510. 10.1038/s41556-019-0299-0

Cheng, H.-F., Wang, C. J., Moeckel, G. W., Zhang, M.-Z., McKanna, J. A., & Harris, R. C. (2002). Cyclooxygenase-2 inhibitor blocks expression of mediators of renal injury in a model of diabetes and hypertension. Kidney International, 62(3), 929–939. 10.1046/j.1523-1755.2002.00520.x

Cui, J., Xu, W., Chen, J., Li, H., Dai, L., Frank, J. A., Peng, S., Wang, S., & Chen, G. (2017). M2 polarization of macrophages facilitates arsenic-induced cell transformation of lung epithelial cells. Oncotarget, 8(13), 21398–21409. 10.18632/oncotarget.15232

Damuzzo, V., Pinton, L., Desantis, G., Solito, S., Marigo, I., Bronte, V., & Mandruzzato, S. (2015). Complexity and challenges in defining myeloid-derived suppressor cells. Cytometry. Part B, Clinical Cytometry, 88(2), 77–91. 10.1002/cyto.b.21206

Dangleben, N. L., Skibola, C. F., & Smith, M. T. (2013). Arsenic immunotoxicity: A review. Environmental Health: A Global Access Science Source, 12(1), 73. 10.1186/1476-069X-12-73

Das, A., Sanyal, T., Bhattacharjee, P., & Bhattacharjee, P. (2021). Depletion of S-adenosylmethionine pool and promoter hypermethylation of *Arsenite methyltransferase* in arsenic-induced skin lesion individuals: A case-control study from West Bengal, India. Environmental Research, 198, 111184. 10.1016/j.envres.2021.111184

Feoktistova, M., Geserick, P., & Leverkus, M. (2016). Crystal Violet Assay for Determining Viability of Cultured Cells. Cold Spring Harbor Protocols, 2016(4), pdb.prot087379. 10.1101/pdb.prot087379

Ferreccio, C., González, C., Milosavjlevic, V., Marshall, G., Sancha, A. M., & Smith, A. H. (2000). Lung cancer and arsenic concentrations in drinking water in Chile. Epidemiology (Cambridge, Mass.), 11(6), 673–679. 10.1097/00001648-200011000-00010

Fiorentino, D. F., Zlotnik, A., Mosmann, T. R., Howard, M., & O’Garra, A. (1991). IL-10 inhibits cytokine production by activated macrophages. Journal of Immunology (Baltimore, Md.: 1950), 147(11), 3815–3822.

Foks, A. C., Van Puijvelde, G. H. M., Wolbert, J., Kröner, M. J., Frodermann, V., Van Der Heijden, T., Van Santbrink, P. J., Boon, L., Bot, I., & Kuiper, J. (2016). CD11b+Gr-1+ myeloid-derived suppressor cells reduce atherosclerotic lesion development in LDLr deficient mice. Cardiovascular Research, 111(3), 252–261. 10.1093/cvr/cvw114

Gamble, M. V., Liu, X., Ahsan, H., Pilsner, R., Ilievski, V., Slavkovich, V., Parvez, F., Levy, D., Factor-Litvak, P., & Graziano, J. H. (2005). Folate, homocysteine, and arsenic metabolism in arsenic-exposed individuals in Bangladesh. Environmental Health Perspectives, 113(12), 1683–1688. 10.1289/ehp.8084

Ge, Z., & Ding, S. (2020). The Crosstalk Between Tumor-Associated Macrophages (TAMs) and Tumor Cells and the Corresponding Targeted Therapy. Frontiers in Oncology, 10. 10.3389/fonc.2020.590941

Giles, B. H., & Mann, K. K. (2022). Arsenic as an immunotoxicant. Toxicology and Applied Pharmacology, 454, 116248. 10.1016/j.taap.2022.116248

Guigni, B. A., van der Velden, J., Kinsey, C. M., Carson, J. A., & Toth, M. J. (2020). Effects of conditioned media from murine lung cancer cells and human tumor cells on cultured myotubes. American Journal of Physiology - Endocrinology and Metabolism, 318(1), E22–E32. 10.1152/ajpendo.00310.2019

Gunawardana, H., Romero, T., Yao, N., Heidt, S., Mulder, A., Elashoff, D. A., & Valenzuela, N. M. (2021). Tissue-specific endothelial cell heterogeneity contributes to unequal inflammatory responses. Scientific Reports, 11(1), 1949. 10.1038/s41598-020-80102-w

Hajishengallis, G., & Russell, M. W. (2015). Innate Humoral Defense Factors. Mucosal Immunology, 251–270. 10.1016/B978-0-12-415847-4.00015-X

Hirayama, D., Iida, T., & Nakase, H. (2017). The Phagocytic Function of Macrophage-Enforcing Innate Immunity and Tissue Homeostasis. International Journal of Molecular Sciences, 19(1), 92. 10.3390/ijms19010092

Hung, C.-C., Zhen, Y.-Y., Niu, S.-W., Hsu, J.-F., Lee, T.-H., Chuang, H.-H., Wang, P.-H., Lee, S.-C., Lin, P.-C., Chiu, Y.-W., Wu, C.-H., Huang, M.-S., Hsiao, M., Chen, H.-C., & Yang, C.-J. (2020). Lung Cancer Cell-Derived Secretome Mediates Paraneoplastic Inflammation and Fibrosis in Kidney in Mice. Cancers, 12(12), 3561. 10.3390/cancers12123561

IARC. (2012). Arsenic, metals, fibres, and dusts (pp. 11–465).

Illingworth, E. J., Rychlik, K. A., Maertens, A., & Sillé, F. C. M. (2025). Sex-specific transcriptomic effects of low-dose inorganic arsenic exposure on bone marrow-derived macrophages. Toxicology, 510, 153988. 10.1016/j.tox.2024.153988

Inaba, T., Sano, H., Kawahito, Y., Hla, T., Akita, K., Toda, M., Yamashina, I., Inoue, M., & Nakada, H. (2003). Induction of cyclooxygenase-2 in monocyte/macrophage by mucins secreted from colon cancer cells. Proceedings of the National Academy of Sciences of the United States of America, 100(5), 2736–2741. 10.1073/pnas.0435410100

Intarasunanont, P., Navasumrit, P., Waraprasit, S., Chaisatra, K., Suk, W. A., Mahidol, C., & Ruchirawat, M. (2012). Effects of arsenic exposure on DNA methylation in cord blood samples from newborn babies and in a human lymphoblast cell line. Environmental Health: A Global Access Science Source, 11, 31. 10.1186/1476-069X-11-31

Jayasingam, S. D., Citartan, M., Thang, T. H., Mat Zin, A. A., Ang, K. C., & Ch’ng, E. S. (2019). Evaluating the Polarization of Tumor-Associated Macrophages Into M1 and M2 Phenotypes in Human Cancer Tissue: Technicalities and Challenges in Routine Clinical Practice. Frontiers in Oncology, 9, 1512. 10.3389/fonc.2019.01512

Jiang, W., Wang, X., Osborne, O. J., Du, Y., Chang, C. H., Liao, Y.-P., Sun, B., Jiang, J., Ji, Z., Li, R., Liu, X., Lu, J., Lin, S., Meng, H., Xia, T., & Nel, A. E. (2017). Pro-Inflammatory and Pro-Fibrogenic Effects of Ionic and Particulate Arsenide and Indium-Containing Semiconductor Materials in the Murine Lung. ACS Nano, 11(2), 1869–1883. 10.1021/acsnano.6b07895

Jiang, X., Wang, J., Deng, X., Xiong, F., Ge, J., Xiang, B., Wu, X., Ma, J., Zhou, M., Li, X., Li, Y., Li, G., Xiong, W., Guo, C., & Zeng, Z. (2019). Role of the tumor microenvironment in PD-L1/PD-1-mediated tumor immune escape. Molecular Cancer, 18(1), 10. 10.1186/s12943-018-0928-4

Kashiwagi, E., Inoue, S., Mizushima, T., Chen, J., Ide, H., Kawahara, T., Reis, L. O., Baras, A. S., Netto, G. J., & Miyamoto, H. (2018). Prostaglandin receptors induce urothelial tumourigenesis as well as bladder cancer progression and cisplatin resistance presumably via modulating PTEN expression. British Journal of Cancer, 118(2), 213–223. 10.1038/bjc.2017.393

Kelly, B., & O’Neill, L. A. J. (2015). Metabolic reprogramming in macrophages and dendritic cells in innate immunity. Cell Research, 25(7), 771–784. 10.1038/cr.2015.68

Klein, S. L., & Flanagan, K. L. (2016). Sex differences in immune responses. Nature Reviews Immunology, 16(10), Article 10. 10.1038/nri.2016.90

Kopp, F., Komatsu, T., Nomura, D. K., Trauger, S. A., Thomas, J. R., Siuzdak, G., Simon, G. M., & Cravatt, B. F. (2010). The Glycerophospho Metabolome and Its Influence on Amino Acid Homeostasis Revealed by Brain Metabolomics of GDE1(−/−) Mice. Chemistry & Biology, 17(8), 831–840. 10.1016/j.chembiol.2010.06.009

Kozul, C. D., Hampton, T. H., Davey, J. C., Gosse, J. A., Nomikos, A. P., Eisenhauer, P. L., Weiss, D. J., Thorpe, J. E., Ihnat, M. A., & Hamilton, J. W. (2009). Chronic exposure to arsenic in the drinking water alters the expression of immune response genes in mouse lung. Environmental Health Perspectives, 117(7), 1108–1115. 10.1289/ehp.0800199

Kzhyshkowska, J., Mamidi, S., Gratchev, A., Kremmer, E., Schmuttermaier, C., Krusell, L., Haus, G., Utikal, J., Schledzewski, K., Scholtze, J., & Goerdt, S. (2006). Novel stabilin-1 interacting chitinase-like protein (SI-CLP) is up-regulated in alternatively activated macrophages and secreted via lysosomal pathway. Blood, 107(8), 3221–3228. 10.1182/blood-2005-07-2843

Lee, C. M., & Hu, J. (2013). Cell density during differentiation can alter the phenotype of bone marrow-derived macrophages. Cell & Bioscience, 3(1), 30. 10.1186/2045-3701-3-30

Lemaire, M., Lemarié, C. A., Molina, M. F., Schiffrin, E. L., Lehoux, S., & Mann, K. K. (2011). Exposure to moderate arsenic concentrations increases atherosclerosis in ApoE-/-mouse model. Toxicological Sciences: An Official Journal of the Society of Toxicology, 122(1), 211–221. 10.1093/toxsci/kfr097

Lemaire, M., Negro Silva, L. F., Lemarié, C. A., Bolt, A. M., Flores Molina, M., Krohn, R. M., Smits, J. E., Lehoux, S., & Mann, K. K. (2015). Arsenic Exposure Increases Monocyte Adhesion to the Vascular Endothelium, a Pro-Atherogenic Mechanism. PloS One, 10(9), e0136592. 10.1371/journal.pone.0136592

Lemarie, A., Bourdonnay, E., Morzadec, C., Fardel, O., & Vernhet, L. (2008). Inorganic arsenic activates reduced NADPH oxidase in human primary macrophages through a Rho kinase/p38 kinase pathway. Journal of Immunology (Baltimore, Md.: 1950), 180(9), 6010–6017. 10.4049/jimmunol.180.9.6010

Lemarie, A., Morzadec, C., Bourdonnay, E., Fardel, O., & Vernhet, L. (2006). Human macrophages constitute targets for immunotoxic inorganic arsenic. Journal of Immunology (Baltimore, Md.: 1950), 177(5), 3019–3027. 10.4049/jimmunol.177.5.3019

Li, L., Wei, K., Ding, Y., Ahati, P., Xu, H., Fang, H., & Wang, H. (2021). M2a Macrophage-Secreted CHI3L1 Promotes Extracellular Matrix Metabolic Imbalances via Activation of IL-13Rα2/MAPK Pathway in Rat Intervertebral Disc Degeneration. Frontiers in Immunology, 12, 666361. 10.3389/fimmu.2021.666361

Lindberg, A.-L., Ekström, E.-C., Nermell, B., Rahman, M., Lönnerdal, B., Persson, L.-A., & Vahter, M. (2008). Gender and age differences in the metabolism of inorganic arsenic in a highly exposed population in Bangladesh. Environmental Research, 106(1), 110–120. 10.1016/j.envres.2007.08.011

Listenberger, L. L., & Brown, D. A. (2007). Fluorescent detection of lipid droplets and associated proteins. Current Protocols in Cell Biology, Chapter 24, Unit 24.2. 10.1002/0471143030.cb2402s35

Liu, H., Dinkova-Kostova, A. T., & Talalay, P. (2008). Coordinate regulation of enzyme markers for inflammation and for protection against oxidants and electrophiles. Proceedings of the National Academy of Sciences of the United States of America, 105(41), 15926–15931. 10.1073/pnas.0808346105

Liu, J., Geng, X., Hou, J., & Wu, G. (2021). New insights into M1/M2 macrophages: Key modulators in cancer progression. Cancer Cell International, 21(1), 389. 10.1186/s12935-021-02089-2

Liu, J., Wu, F., & Zhou, H. (2020). Macrophage-derived exosomes in cancers: Biogenesis, functions and therapeutic applications. Immunology Letters, 227, 102–108. 10.1016/j.imlet.2020.08.003

Liu, Y., Zugazagoitia, J., Ahmed, F. S., Henick, B. S., Gettinger, S. N., Herbst, R. S., Schalper, K. A., & Rimm, D. L. (2020). Immune Cell PD-L1 Colocalizes with Macrophages and Is Associated with Outcome in PD-1 Pathway Blockade Therapy. Clinical Cancer Research: An Official Journal of the American Association for Cancer Research, 26(4), 970–977. 10.1158/1078-0432.CCR-19-1040

Liu, Z., Hou, Y., Li, L., Yang, Y., Jia, J., Hong, Z., Li, T., Xu, Y., Fu, J., Sun, Y., Yamamoto, M., Wang, H., & Pi, J. (2019). Nrf2 deficiency aggravates the increase in osteoclastogenesis and bone loss induced by inorganic arsenic. Toxicology and Applied Pharmacology, 367, 62–70. 10.1016/j.taap.2019.02.003

Livak, K. J., & Schmittgen, T. D. (2001). Analysis of relative gene expression data using real-time quantitative PCR and the 2(-Delta Delta C(T)) Method. Methods (San Diego, Calif.), 25(4), 402–408. 10.1006/meth.2001.1262

Mantovani, A., Sica, A., Sozzani, S., Allavena, P., Vecchi, A., & Locati, M. (2004). The chemokine system in diverse forms of macrophage activation and polarization. Trends in Immunology, 25(12), 677–686. 10.1016/j.it.2004.09.015

Martinez, F. O., & Gordon, S. (2014). The M1 and M2 paradigm of macrophage activation: Time for reassessment. F1000prime Reports, 6, 13. 10.12703/P6-13

Medina, S., Lauer, F. T., Castillo, E. F., Bolt, A. M., Ali, A.-M. S., Liu, K. J., & Burchiel, S. W. (2020). Exposures to uranium and arsenic alter intraepithelial and innate immune cells in the small intestine of male and female mice. Toxicology and Applied Pharmacology, 403, 115155. 10.1016/j.taap.2020.115155

Michel-Ramirez, G., Recio-Vega, R., Lantz, R. C., Gandolfi, A. J., Olivas-Calderon, E., Chau, B. T., & Amistadi, M. K. (2020). Assessment of YAP gene polymorphisms and arsenic interaction in Mexican women with breast cancer. Journal of Applied Toxicology: JAT, 40(3), 342–351. 10.1002/jat.3907

Mills, C. D. (2001). Macrophage arginine metabolism to ornithine/urea or nitric oxide/citrulline: A life or death issue. Critical Reviews in Immunology, 21(5), 399– 425.

Mills, C. D., Kincaid, K., Alt, J. M., Heilman, M. J., & Hill, A. M. (2000). M-1/M-2 macrophages and the Th1/Th2 paradigm. Journal of Immunology (Baltimore, Md.: 1950), 164(12), 6166–6173. 10.4049/jimmunol.164.12.6166

Mollace, V., Muscoli, C., Masini, E., Cuzzocrea, S., & Salvemini, D. (2005). Modulation of prostaglandin biosynthesis by nitric oxide and nitric oxide donors. Pharmacological Reviews, 57(2), 217–252. 10.1124/pr.57.2.1

Morgan, P. K., Huynh, K., Pernes, G., Miotto, P. M., Mellett, N. A., Giles, C., Meikle, P. J., Murphy, A. J., & Lancaster, G. I. (2021). Macrophage polarization state affects lipid composition and the channeling of exogenous fatty acids into endogenous lipid pools. The Journal of Biological Chemistry, 297(6), 101341. 10.1016/j.jbc.2021.101341

Mosser, D. M., & Edwards, J. P. (2008). Exploring the full spectrum of macrophage activation. Nature Reviews. Immunology, 8(12), 958–969. 10.1038/nri2448

Muhetaer, M., Yang, M., Xia, R., Lai, Y., & Wu, J. (2022). Gender difference in arsenic biotransformation is an important metabolic basis for arsenic toxicity. BMC Pharmacology and Toxicology, 23(1), 15. 10.1186/s40360-022-00554-w

Nath, N., & Kashfi, K. (2020). Tumor associated macrophages and ‘NO.’ Biochemical Pharmacology, 176, 113899. 10.1016/j.bcp.2020.113899

Negro Silva, L. F., Lemaire, M., Lemarié, C. A., Plourde, D., Bolt, A. M., Chiavatti, C., Bohle, D. S., Slavkovich, V., Graziano, J. H., Lehoux, S., & Mann, K. K. (2017). Effects of Inorganic Arsenic, Methylated Arsenicals, and Arsenobetaine on Atherosclerosis in the Mouse Model and the Role of As3mt-Mediated Methylation. Environmental Health Perspectives, 125(7), 077001. 10.1289/EHP806

Niedzwiecki, M. M., Hall, M. N., Liu, X., Oka, J., Harper, K. N., Slavkovich, V., Ilievski, V., Levy, D., van Geen, A., Mey, J. L., Alam, S., Siddique, A. B., Parvez, F., Graziano, J. H., & Gamble, M. V. (2013). A dose-response study of arsenic exposure and global methylation of peripheral blood mononuclear cell DNA in Bangladeshi adults. Environmental Health Perspectives, 121(11–12), 1306–1312. 10.1289/ehp.1206421

Nishikawa, M., Kurano, M., Ikeda, H., Aoki, J., & Yatomi, Y. (2015). Lysophosphatidylserine has Bilateral Effects on Macrophages in the Pathogenesis of Atherosclerosis. Journal of Atherosclerosis and Thrombosis, 22(5), 518–526. 10.5551/jat.25650

Okoji, R. S., Yu, R. C., Maronpot, R. R., & Froines, J. R. (2002). Sodium arsenite administration via drinking water increases genome-wide and Ha-ras DNA hypomethylation in methyl-deficient C57BL/6J mice. Carcinogenesis, 23(5), 777– 785. 10.1093/carcin/23.5.777

Ooki, A., Del Carmen Rodriguez Pena, M., Marchionni, L., Dinalankara, W., Begum, A., Hahn, N. M., VandenBussche, C. J., Rasheed, Z. A., Mao, S., Netto, G. J., Sidransky, D., & Hoque, M. O. (2018). YAP1 and COX2 Coordinately Regulate Urothelial Cancer Stem-like Cells. Cancer Research, 78(1), 168–181. 10.1158/0008-5472.CAN-17-0836

Orecchioni, M., Ghosheh, Y., Pramod, A. B., & Ley, K. (2019). Macrophage Polarization: Different Gene Signatures in M1(LPS+) vs. Classically and M2(LPS-) vs. Alternatively Activated Macrophages. Frontiers in Immunology, 10, 1084. 10.3389/fimmu.2019.01084

Parameswaran, N., & Patial, S. (2010). Tumor necrosis factor-α signaling in macrophages. Critical Reviews in Eukaryotic Gene Expression, 20(2), 87–103. 10.1615/critreveukargeneexpr.v20.i2.10

Parvez, F., Chen, Y., Yunus, M., Olopade, C., Segers, S., Slavkovich, V., Argos, M., Hasan, R., Ahmed, A., Islam, T., Akter, M. M., Graziano, J. H., & Ahsan, H. (2013). Arsenic exposure and impaired lung function. Findings from a large population-based prospective cohort study. American Journal of Respiratory and Critical Care Medicine, 188(7), 813–819. 10.1164/rccm.201212-2282OC

Percie du Sert, N., Hurst, V., Ahluwalia, A., Alam, S., Avey, M. T., Baker, M., Browne, W. J., Clark, A., Cuthill, I. C., Dirnagl, U., Emerson, M., Garner, P., Holgate, S. T., Howells, D. W., Karp, N. A., Lazic, S. E., Lidster, K., MacCallum, C. J., Macleod, M., … Würbel, H. (2020). The ARRIVE guidelines 2.0: Updated guidelines for reporting animal research. BMC Veterinary Research, 16(1), 242. 10.1186/s12917-020-02451-y

Pierce, B. L., Tong, L., Argos, M., Gao, J., Jasmine, F., Roy, S., Paul-Brutus, R., Rahaman, R., Rakibuz-Zaman, M., Parvez, F., Ahmed, A., Quasem, I., Hore, S. K., Alam, S., Islam, T., Harjes, J., Sarwar, G., Slavkovich, V., Gamble, M. V., … Ahsan, H. (2013). Arsenic metabolism efficiency has a causal role in arsenic toxicity: Mendelian randomization and gene-environment interaction. International Journal of Epidemiology, 42(6), 1862–1872. 10.1093/ije/dyt182

Pitman, M. R., Lewis, A. C., Davies, L. T., Moretti, P. A. B., Anderson, D., Creek, D. J., Powell, J. A., & Pitson, S. M. (2022). The sphingosine 1-phosphate receptor 2/4 antagonist JTE-013 elicits off-target effects on sphingolipid metabolism. Scientific Reports, 12(1), 454. 10.1038/s41598-021-04009-w

Podgorski, J., & Berg, M. (2020). Global threat of arsenic in groundwater. Science (New York, N.Y.), 368(6493), 845–850. 10.1126/science.aba1510

Poledne, R., Malinska, H., Kubatova, H., Fronek, J., Thieme, F., Kauerova, S., & Lesna, I. K. (2019). Polarization of Macrophages in Human Adipose Tissue is Related to the Fatty Acid Spectrum in Membrane Phospholipids. Nutrients, 12(1), 8. 10.3390/nu12010008

Quandt, J., & Dorovini-Zis, K. (2004). The beta chemokines CCL4 and CCL5 enhance adhesion of specific CD4+ T cell subsets to human brain endothelial cells. Journal of Neuropathology and Experimental Neurology, 63(4), 350–362. 10.1093/jnen/63.4.350

Rabold, K., Aschenbrenner, A., Thiele, C., Boahen, C. K., Schiltmans, A., Smit, J. W. A., Schultze, J. L., Netea, M. G., Adema, G. J., & Netea-Maier, R. T. (2020). Enhanced lipid biosynthesis in human tumor-induced macrophages contributes to their protumoral characteristics. Journal for Immunotherapy of Cancer, 8(2), e000638. 10.1136/jitc-2020-000638

Rahman, A., Vahter, M., Ekström, E.-C., & Persson, L.-Å. (2011). Arsenic exposure in pregnancy increases the risk of lower respiratory tract infection and diarrhea during infancy in Bangladesh. Environmental Health Perspectives, 119(5), 719–724. 10.1289/ehp.1002265

Rahman, F. A., Angus, S. A., Stokes, K., Karpowicz, P., & Krause, M. P. (2020). Impaired ECM Remodeling and Macrophage Activity Define Necrosis and Regeneration Following Damage in Aged Skeletal Muscle. International Journal of Molecular Sciences, 21(13), 4575. 10.3390/ijms21134575

Rahman, M., Vahter, M., Sohel, N., Yunus, M., Wahed, M. A., Streatfield, P. K., Ekström, E.-C., & Persson, L. A. (2006). Arsenic exposure and age and sex-specific risk for skin lesions: A population-based case-referent study in Bangladesh. Environmental Health Perspectives, 114(12), 1847–1852. 10.1289/ehp.9207

Raqib, R., Ahmed, S., Ahsan, K. B., Kippler, M., Akhtar, E., Roy, A. K., Lu, Y., Arifeen, S. E., Wagatsuma, Y., & Vahter, M. (2017). Humoral Immunity in Arsenic-Exposed Children in Rural Bangladesh: Total Immunoglobulins and Vaccine-Specific Antibodies. Environmental Health Perspectives, 125(6), 067006. 10.1289/EHP318

Raqib, R., Akhtar, E., Haq, Md. A., & Sarker, P. (2023). Effects of prenatal exposure to arsenic on T cell development in children. Current Opinion in Toxicology, 34, 100389. 10.1016/j.cotox.2023.100389

Ren, W., Xia, Y., Chen, S., Wu, G., Bazer, F. W., Zhou, B., Tan, B., Zhu, G., Deng, J., & Yin, Y. (2019). Glutamine Metabolism in Macrophages: A Novel Target for Obesity/Type 2 Diabetes. Advances in Nutrition (Bethesda, Md.), 10(2), 321–330. 10.1093/advances/nmy084

Ribeiro, S. P., Villar, J., Downey, G. P., Edelson, J. D., & Slutsky, A. S. (1996). Effects of the stress response in septic rats and LPS-stimulated alveolar macrophages: Evidence for TNF-alpha posttranslational regulation. American Journal of Respiratory and Critical Care Medicine, 154(6 Pt 1), 1843–1850. 10.1164/ajrccm.154.6.8970379

Rodríguez, J. P., Casas, J., Balboa, M. A., & Balsinde, J. (2025). Bioactive lipid signaling and lipidomics in macrophage polarization: Impact on inflammation and immune regulation. Frontiers in Immunology, 16, 1550500. 10.3389/fimmu.2025.1550500

Rosenblat, M., Oren, R., & Aviram, M. (2006). Lysophosphatidylcholine (LPC) attenuates macrophage-mediated oxidation of LDL. Biochemical and Biophysical Research Communications, 344(4), 1271–1277. 10.1016/j.bbrc.2006.04.038

Rostam, H. M., Reynolds, P. M., Alexander, M. R., Gadegaard, N., & Ghaemmaghami, A. M. (2017). Image based Machine Learning for identification of macrophage subsets. Scientific Reports, 7(1), 3521. 10.1038/s41598-017-03780-z

Rőszer, T. (2015). Understanding the Mysterious M2 Macrophage through Activation Markers and Effector Mechanisms. Mediators of Inflammation, 2015, 816460. 10.1155/2015/816460

Sadiku, P., & Walmsley, S. R. (2019). Hypoxia and the regulation of myeloid cell metabolic imprinting: Consequences for the inflammatory response. EMBO Reports, 20(5), e47388. 10.15252/embr.201847388

Sakurai, T., Kaise, T., & Matsubara, C. (1998). Inorganic and methylated arsenic compounds induce cell death in murine macrophages via different mechanisms. Chemical Research in Toxicology, 11(4), 273–283. 10.1021/tx9701384

Sakurai, T., Ohta, T., Tomita, N., Kojima, C., Hariya, Y., Mizukami, A., & Fujiwara, K. (2006). Evaluation of immunotoxic and immunodisruptive effects of inorganic arsenite on human monocytes/macrophages. International Immunopharmacology, 6(2), 304–315. 10.1016/j.intimp.2005.06.012

Saleh, L. S., Vanderheyden, C., Frederickson, A., & Bryant, S. J. (2020). Prostaglandin E2 and Its Receptor EP2 Modulate Macrophage Activation and Fusion in Vitro. ACS Biomaterials Science & Engineering, 6(5), 2668–2681. 10.1021/acsbiomaterials.9b01180

Salvemini, D., Kim, S. F., & Mollace, V. (2013). Reciprocal regulation of the nitric oxide and cyclooxygenase pathway in pathophysiology: Relevance and clinical implications. American Journal of Physiology. Regulatory, Integrative and Comparative Physiology, 304(7), R473–487. 10.1152/ajpregu.00355.2012

Schurz, H., Salie, M., Tromp, G., Hoal, E. G., Kinnear, C. J., & Möller, M. (2019). The X chromosome and sex-specific effects in infectious disease susceptibility. Human Genomics, 13(1), 2. 10.1186/s40246-018-0185-z

Sengupta, M., & Bishayi, B. (2002). Effect of lead and arsenic on murine macrophage response. Drug and Chemical Toxicology, 25(4), 459–472. 10.1081/dct-120014796

Serbulea, V., Upchurch, C. M., Schappe, M. S., Voigt, P., DeWeese, D. E., Desai, B. N., Meher, A. K., & Leitinger, N. (2018). Macrophage phenotype and bioenergetics are controlled by oxidized phospholipids identified in lean and obese adipose tissue. Proceedings of the National Academy of Sciences of the United States of America, 115(27), E6254–E6263. 10.1073/pnas.1800544115

Shaji, E., Santosh, M., Sarath, K. V., Prakash, P., Deepchand, V., & Divya, B. V. (2021). Arsenic contamination of groundwater: A global synopsis with focus on the Indian Peninsula. Geoscience Frontiers, 12(3), 101079. 10.1016/j.gsf.2020.08.015

Shang, Y., Zhu, T., Lenz, A.-G., Frankenberger, B., Tian, F., Chen, C., & Stoeger, T. (2013). Reduced in vitro toxicity of fine particulate matter collected during the 2008 Summer Olympic Games in Beijing: The roles of chemical and biological components. Toxicology in Vitro: An International Journal Published in Association with BIBRA, 27(7), 2084–2093. 10.1016/j.tiv.2013.08.004

Smith, A. H., Marshall, G., Yuan, Y., Ferreccio, C., Liaw, J., von Ehrenstein, O., Steinmaus, C., Bates, M. N., & Selvin, S. (2006). Increased mortality from lung cancer and bronchiectasis in young adults after exposure to arsenic in utero and in early childhood. Environmental Health Perspectives, 114(8), 1293–1296. 10.1289/ehp.8832

Smith, A. H., Marshall, G., Yuan, Y., Liaw, J., Ferreccio, C., & Steinmaus, C. (2011). Evidence from Chile that arsenic in drinking water may increase mortality from pulmonary tuberculosis. American Journal of Epidemiology, 173(4), 414–420. 10.1093/aje/kwq383

Syrett, C. M., Sindhava, V., Sierra, I., Dubin, A. H., Atchison, M., & Anguera, M. C. (2019). Diversity of Epigenetic Features of the Inactive X-Chromosome in NK Cells, Dendritic Cells, and Macrophages. Frontiers in Immunology, 9. 10.3389/fimmu.2018.03087

Schmidt, T., Kahn, R., Kahn, F. (2022). Ascorbic acid attenuates activation and cytokine production in sepsis-like monocytes. Journal of Leukocyte Biology, 112(3). 10.1002/JLB.4AB0521-243R

Thompson, B. J. (2020). YAP/TAZ: Drivers of Tumor Growth, Metastasis, and Resistance to Therapy. BioEssays: News and Reviews in Molecular, Cellular and Developmental Biology, 42(5), e1900162. 10.1002/bies.201900162

Tokar, E. J., Diwan, B. A., & Waalkes, M. P. (2010). Arsenic Exposure Transforms Human Epithelial Stem/Progenitor Cells into a Cancer Stem-like Phenotype. Environmental Health Perspectives, 118(1), 108–115. 10.1289/ehp.0901059

Trauelsen, M., Hiron, T. K., Lin, D., Petersen, J. E., Breton, B., Husted, A. S., Hjorth, S. A., Inoue, A., Frimurer, T. M., Bouvier, M., O’Callaghan, C. A., & Schwartz, T. W. (2021). Extracellular succinate hyperpolarizes M2 macrophages through SUCNR1/GPR91-mediated Gq signaling. Cell Reports, 35(11), 109246. 10.1016/j.celrep.2021.109246

Tripathi, P. (2007). Nitric oxide and immune response. Indian Journal of Biochemistry & Biophysics, 44(5), 310–319.

Valentin, J. E., Stewart-Akers, A. M., Gilbert, T. W., & Badylak, S. F. (2009). Macrophage Participation in the Degradation and Remodeling of Extracellular Matrix Scaffolds. Tissue Engineering. Part A, 15(7), 1687–1694. 10.1089/ten.tea.2008.0419

Veglia, F., Sanseviero, E., & Gabrilovich, D. I. (2021). Myeloid-derived suppressor cells in the era of increasing myeloid cell diversity. Nature Reviews. Immunology, 21(8), 485–498. 10.1038/s41577-020-00490-y

Versteeg, L., Le Guezennec, X., Zhan, B., Liu, Z., Angagaw, M., Woodhouse, J. D., Biswas, S., & Beaumier, C. M. (2017). Transferring Luminex® cytokine assays to a wall-less plate technology: Validation and comparison study with plasma and cell culture supernatants. Journal of Immunological Methods, 440, 74–82. 10.1016/j.jim.2016.11.003

Wang, C., Ma, C., Gong, L., Guo, Y., Fu, K., Zhang, Y., Zhou, H., & Li, Y. (2021). Macrophage Polarization and Its Role in Liver Disease. Frontiers in Immunology, 12, 803037. 10.3389/fimmu.2021.803037

Weigert, A., Olesch, C., & Brüne, B. (2019). Sphingosine-1-Phosphate and Macrophage Biology-How the Sphinx Tames the Big Eater. Frontiers in Immunology, 10, 1706. 10.3389/fimmu.2019.01706

WHO. (2006). Protecting groundwater for health Managing the quality of drinking-water sources. https://www.who.int/publications/i/item/9241546689

Winkler, J., Abisoye-Ogunniyan, A., Metcalf, K. J., & Werb, Z. (2020). Concepts of extracellular matrix remodelling in tumour progression and metastasis. Nature Communications, 11(1), 5120. 10.1038/s41467-020-18794-x

Xue, J., Xiao, T., Wei, S., Sun, J., Zou, Z., Shi, M., Sun, Q., Dai, X., Wu, L., Li, J., Xia, H., Tang, H., Zhang, A., & Liu, Q. (2021). miR-21-regulated M2 polarization of macrophage is involved in arsenicosis-induced hepatic fibrosis through the activation of hepatic stellate cells. Journal of Cellular Physiology, 236(8), 6025– 6041. 10.1002/jcp.30288

Yang, J., Fan, Y., Xie, B., & Yang, D. (2021). A Combination of RNA-Seq Analysis and Use of TCGA Database for Determining the Molecular Mechanism and Identifying Potential Drugs for GJB1 in Ovarian Cancer. OncoTargets and Therapy, 14, 2623–2633. 10.2147/OTT.S303589

Yang, Y., Li, C., Liu, T., Dai, X., & Bazhin, A. V. (2020). Myeloid-Derived Suppressor Cells in Tumors: From Mechanisms to Antigen Specificity and Microenvironmental Regulation. Frontiers in Immunology, 11, 1371. 10.3389/fimmu.2020.01371

Zajd, C. M., Ziemba, A. M., Miralles, G. M., Nguyen, T., Feustel, P. J., Dunn, S. M., Gilbert, R. J., & Lennartz, M. R. (2020). Bone Marrow-Derived and Elicited Peritoneal Macrophages Are Not Created Equal: The Questions Asked Dictate the Cell Type Used. Frontiers in Immunology, 11, 269. 10.3389/fimmu.2020.00269

Zaravinos, A., Lambrou, G. I., Boulalas, I., Delakas, D., & Spandidos, D. A. (2011). Identification of common differentially expressed genes in urinary bladder cancer. PloS One, 6(4), e18135. 10.1371/journal.pone.0018135

Zhang, F., Wang, H., Wang, X., Jiang, G., Liu, H., Zhang, G., Wang, H., Fang, R., Bu, X., Cai, S., & Du, J. (2016). TGF-β induces M2-like macrophage polarization via SNAIL-mediated suppression of a pro-inflammatory phenotype. Oncotarget, 7(32), 52294–52306. 10.18632/oncotarget.10561

Zhang, M., Qi, Y., Li, H., Cui, J., Dai, L., Frank, J. A., Chen, J., Xu, W., & Chen, G. (2016). AIM2 inflammasome mediates Arsenic-induced secretion of IL-1 β and IL-18. Oncoimmunology, 5(6), e1160182. 10.1080/2162402X.2016.1160182

Zhao, T., Su, Z., Li, Y., Zhang, X., & You, Q. (2020). Chitinase-3 like-protein-1 function and its role in diseases. Signal Transduction and Targeted Therapy, 5(1), 1–20. 10.1038/s41392-020-00303-7

Zhou, X., Liu, X., & Huang, L. (2021). Macrophage-Mediated Tumor Cell Phagocytosis: Opportunity for Nanomedicine Intervention. Advanced Functional Materials, 31(5), 2006220. 10.1002/adfm.202006220

